# Bifurcation Structure and Cross Nuclei Universality Govern Frequency-Selective Deep Brain Stimulation

**DOI:** 10.64898/2026.07.09.737626

**Authors:** Xiangyu Samuel Ma, Milad Lankarany

## Abstract

High-frequency deep brain stimulation (DBS, >90 Hz) reliably suppresses Parkinsonian motor symptoms, whereas sub-therapeutic frequencies (<60 Hz) worsen them, yet the circuit mechanism underlying this frequency selectivity remains unresolved. We develop an analytically tractable excitatory–inhibitory continuous attractor neural network with threshold-linear transfer functions. The model predicts a *boundary equilibrium bifurcation* (BEB) at a critical DBS frequency *f*_th_ that simultaneously accounts for two observations that prior models have treated separately: abrupt excitatory suppression and a concurrent linear rise in GABAergic output with stimulation rate. A separate, orthogonal bifurcation condition — the Hopf boundary 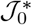— governs endogenous beta oscillations independently of stimulation frequency; DBS suppresses these oscillations by destroying the active fixed point via the BEB, not by crossing the Hopf boundary. The predicted signature — an oscillation frequency that remains at *f*_0_ up to the suppression threshold, then drops discontinuously — is testable with existing intraoperative protocols. The sub-critical spectral sharpness index *C*= 2*f*_0_*/*Δ*f*_FWHM_ diverges at the Hopf boundary and provides a parameter-free biomarker of pathological synchrony computable from any local field potential recording. Fitting the model to intraoperative VIM single-unit recordings across nine stimulation frequencies yields physiologically plausible parameters consistent with the observed transient and steady-state dynamics. Frequency–response data from four DBS nuclei (STN, SNr, VIM, RT) each conform to a rectified quadratic derived from the mean-field equations; nucleus pairs sharing circuit role collapse onto common master curves (*R*^2^ ≥ 0.988 per nucleus, collapsed *R*^2^ ≥ 0.988) after a two-parameter rescaling, with no further free parameters.

**Significance:** High-frequency deep brain stimulation (DBS) relieves Parkinson’s disease motor symptoms, while sub-therapeutic frequencies worsen them; the circuit origin of this ∼90 Hz threshold has lacked explanation. We show that one bifurcation — a boundary equilibrium bifurcation (BEB) at a critical frequency f_th_ — jointly explains the abrupt loss of excitatory firing and the concurrent rise in GABAergic output. The Hopf boundary governing endogenous beta oscillations is independent of stimulation frequency, so DBS suppresses beta by destroying the active state through the BEB, not by shifting the Hopf boundary: disease progression and therapy act on separate axes. Frequency–response curves from four DBS nuclei collapse onto two universality-class master curves after a two-parameter rescaling, yielding cross-nucleus predictions without further fitting.

---

Deep brain stimulation (DBS) is an established therapy for Parkinson’s disease (PD), essential tremor, and dystonia (1, 2). Its efficacy is strongly frequency-dependent: stimulation above approximately 90 Hz (at a fixed inter-pulse interval) consistently improves bradykinesia and rigidity, whereas frequencies below 60 Hz can worsen symptoms or leave them unchanged. This threshold is consistent across DBS targets and patient populations, with parallel frequency-dependence reported in animal models (1, 2), and the same frequency dependence is reflected in the patterned discharge of individual subthalamic and thalamic neurons recorded during intraoperative stimulation (3). The network-level mechanism that makes ~90 Hz a biologically meaningful boundary for improving versus worsening motor symptoms has, however, remained unidentified.

A hallmark of PD pathophysiology is excessive beta-band (13–30 Hz) synchronization in basal ganglia–thalamocortical circuits; its attenuation by high-frequency DBS and by dopaminergic medication correlates with symptomatic improvement (4, 5). At the synaptic level, recent work suggests that high-frequency stimulation preferentially depletes glutamatergic vesicle pools through short-term synaptic depression, thereby shifting the effective excitation–inhibition balance in a frequency-dependent manner (6). Intrinsic neural diversity within a nucleus further shapes how individual neurons respond to this shift (7). Despite this evidence, no quantitative mechanistic account of the 90 Hz threshold had been proposed.

Biophysically detailed STN–GPe conductance-based models demonstrate high-frequency regularization pathological bursting (8), but their tractability is limited and they generate no cross nuclei predictions. Wilson–Cowan E–I frameworks support Hopf bifurcations (9, 10 yet sigmoid transfer functions preclude the boundary equilibrium bifurcation (BEB) that a threshold-linear nonlinearity introduces; the abrupt frequency threshold is therefore absent in those models. Rate-coding desynchronization models (11) target patterned stimulation rather than tonic high-frequency DBS; random-connectivity mean-field extensions (12, 13) produce realistic spectral exponents but not the cross nuclei universality structure. The connection between spectral sharpening and proximity to a dynamical phase transition is a natural expectation from critical slowing-down near such transitions, but has not been derived from a mechanistic DBS circuit model No existing framework simultaneously accounts for the frequency threshold, beta suppression, and multi-nuclear response organization in closed analytical form.

Here we extend the continuous attractor neural network (CANN) framework (14, 15) with bidirectional E–I coupling, ReLU transfer functions, and an exponential spatial connectivity kernel, following the standard CANN construction. The resulting equations admit closed-form expressions for the BEB frequency threshold, endogenous beta oscillations and their DBS-induced suppression, and spatially localized activity profiles. Fitting the model to intraoperative single-unit recordings from nine DBS frequencies yields physiologically plausible parameters; the frequency–response curves across all four nuclei (STN, SNr, VIM, RT) organize into two cross nuclei universality classes, and the corresponding master curves follow from first principles without additional free parameters.

## Results

### Cross nuclei universality of DBS frequency–response

Single-unit recordings obtained during intraoperative DBS at nine frequencies (1–100 Hz) across STN, SNr, VIM, and RT (3) reveal a regularity that the theory accounts for quantitatively.

Across all four nuclei, the mean firing rate as a function of stimulation frequency is consistent with a rectified quadratic 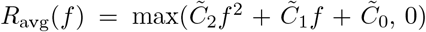, with individual fits *R*^2^ ≥ 0.988 (Fig. 1A,B,D,E; STN 0.989, SNr 0.988, VIM 0.996, RT 0.995). This functional form is not assumed but is a direct consequence of the E–I mean-field equations: substituting the frequency-modulated DBS drive *I*_DBS_(*f*) ∝ *f* (1 + *βf*) into the fixed-point equations yields *R*_*_(*f*) = *C*_0_ + *C*_1_*f* + *C*_2_*f* ^2^ on the high-activity branch, with *C*_2_ ∝*β* setting the concavity (SI Appendix, Section A.4). The parameter *β* reflects the degree of glutamatergic short-term depression: negative values produce a downward concavity that generates the observed therapeutic suppression, while *β* = 0 recovers a linear frequency–response.

**Fig. 1.**
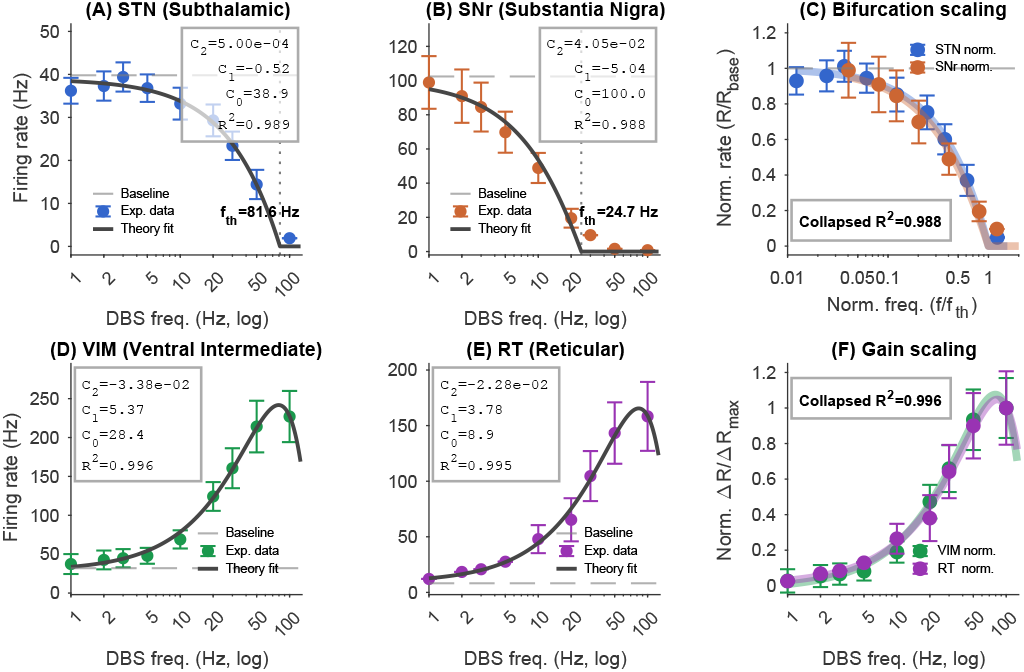
Quadratic frequency–response and cross nuclei data collapse across four DBS targets. **(A)** STN frequency–response. Filled circles: mean firing rates (±SE) from Milosevic et al. (3). Black curve: rectified quadratic fit (*R*^2^ = 0.989); vertical dotted line: fitted threshold 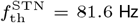. **(B)** SNr frequency–response from the same dataset (*R*^2^ = 0 .988); fitted threshold 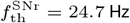. **(C)** *Bifurcation scaling collapse*. STN (blue) and SNr (orange) re-plotted after rescaling frequency by 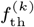 and firing rate by 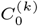. Collapsed *R*^2^ = 0.988. **(D)** VIM frequency–response from the same dataset (*R*^2^ = 0.996). The non-monotonic dip at 30 Hz is discussed in the text. **(E)** RT frequency–response from the same dataset (*R*^2^ = 0.995). **(F)** *Gain scaling collapse*. VIM (green) and RT (purple) after subtracting baseline 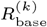 and dividing by 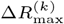. Collapsed *R* ^2^ = 0 .996. Each nucleus fit uses three free parameters 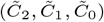; the cross nuclei predictions use no additional parameters.

The four nuclei divide naturally into two classes according to whether their frequency–response is terminated by a threshold (STN, SNr) or is monotonically increasing (VIM, RT). The model predicts that within each class, nuclei should collapse onto a single dimensionless master curve after nucleus-specific rescaling. For STN and SNr (*bifurcation scaling*), dividing frequency by each nucleus’s BEB threshold 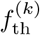 and firing rate by its baseline 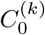 yields 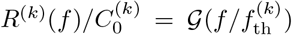; the two curves superimpose with collapsed *R*^2^ = 0.988 (Fig. 1C). For VIM and RT (*gain scaling*), normalizing by the maximum evoked rate change yields 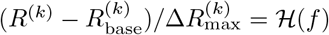; collapsed *R* 2 = 0. 996 (Fig. 1F). The two pairs thus delineate distinct response topologies: threshold-terminated suppression (STN, SNr), in which excitatory firing is suppressed above *f*_th_, versus monotone gain modulation (VIM, RT), in which firing increases with frequency across the stimulation range tested (1–100 Hz).

The two rescaling parameters per nucleus pair have direct biological interpretations. For the bifurcation class, the amplitude ratio 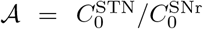reflects the relative baseline excitatory drives of the two nuclei, and the frequency ratio 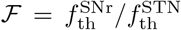 reflects their relative synaptic time constants and depression coefficients through Equation (1). Once the full frequency–response of one nucleus in a pair is measured, the partner’s response at any unstimulated frequency follows without additional parameters: 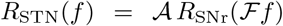) (SI Appendix, Section A.4). Similarly, *R*_VIM_(*f*) = P *R*_RT_(*f*) + Q, with 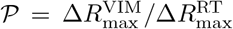. The predictive content of these relations is testable by withholding one or more frequencies from the fit and checking whether the collapsed curve predicts the held-out data.

### Frequency-selective suppression via a boundary equilibrium bifurcation

The cross nuclei organization described above has a specific dynamical origin. In the spatially uniform regime, the E–I network admits two coexisting steady states: a low-activity branch (*R* = *R*^′^ = 0) and a high-activity branch with steady-state rates *R*_*_(*f*) and 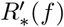 (SI Appendix, Section A.3). As stimulation frequency increases, the depression factor *F* (*f*) = 1 + *βf* (*β <* 0) reduces the effective excitatory drive, lowering the active fixed point toward threshold. The critical frequency *f*_th_ at which the excitatory steady state meets the threshold satisfies:

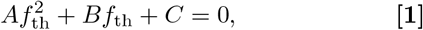

with 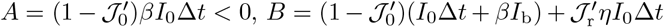 and 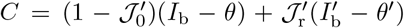 Appendix, Section A.5). This is a boundary equilibrium bifurcation (BEB): unlike a saddle-node bifurcation, the fixed point is not annihilated by collision with another equilibrium but by reaching the non-smooth boundary of the ReLU nonlinearity at *R* = 0. The distinction has quantitative consequences. In a saddle-node bifurcation th<u>e eigenv</u>alue of the vanishing fixed point scales as 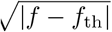, so the relaxation time diverges and the suppression is gradual. In the BEB, the eigenvalues *λ*_1,2_ at the active fixed point remain finite and unchanged right up to *f*_th_; there is no divergence of the relaxation timescale. The suppression is therefore discontinuous in *f* at the level of the mean-field fixed point, a sharp signature that distinguishes the BEB from the gradual, square-root closure characteristic of a saddle-node bifurcation.

Above *f*_th_, the excitatory population is fully suppressed and the inhibitory equation decouples. Above a second threshold 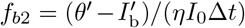, which exceeds *f*_th_ when 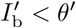, the inhibitory rate grows linearly as:

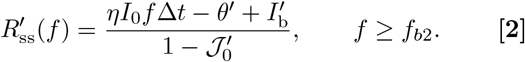

The model thus jointly predicts the therapeutic suppression threshold and the DBS-induced increase in inhibitory output reported by Xu et al. (16), deriving both from the same two equations. Figure 2B confirms Equation (2) against simulation; note that the interval [*f*_th_, *f*_*b*2_] is a silent epoch in which both populations are subthreshold, and the inhibitory rise begins at *f*_*b*2_, not at *f*_th_.

**Fig. 2.**
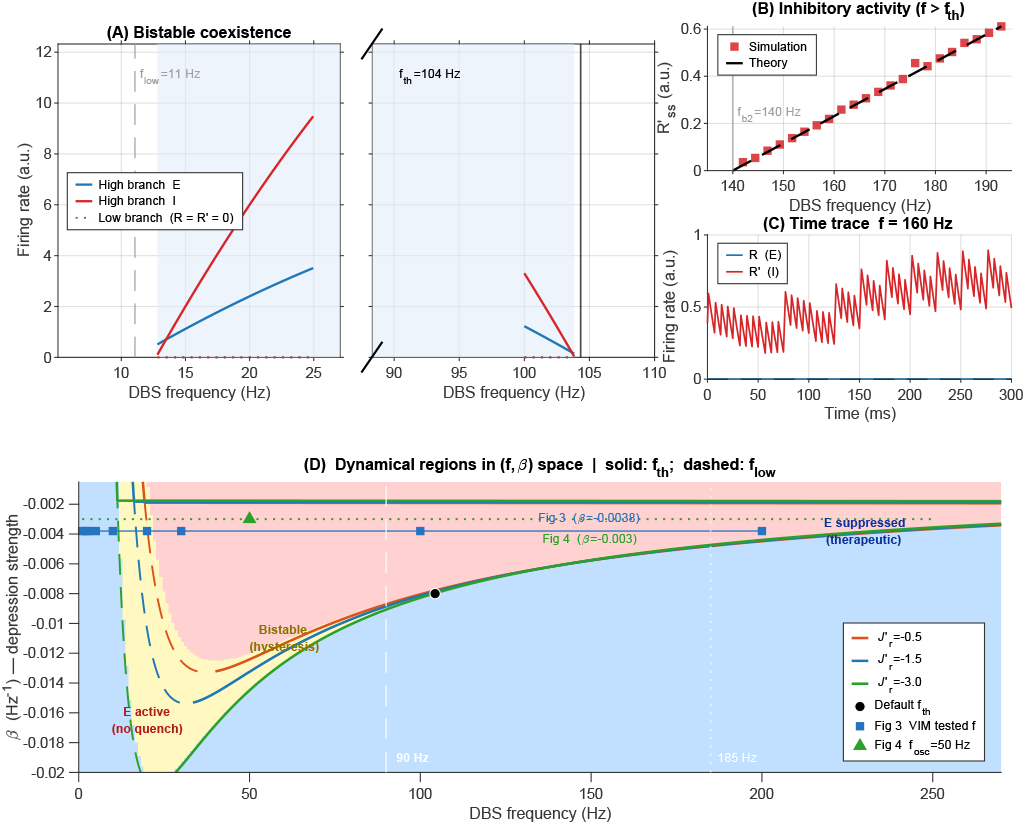
Bistability and boundary equilibrium bifurcation. **(A)** Analytical steady-state firing-rate curves (broken *x*-axis isolating both bifurcation boundaries). Left panel: bistable onset at *f*_low_ ≈ 11 Hz. Right panel: BEB at *f*_th_ ≈ 104 Hz (solid vertical line) where the high-activity branch terminates. Blue: excitatory high-activity branch *R*^high^(*f*); red: inhibitory *R*^*′*high^(*f*); dotted: low-activity branch (*R* = *R*^*′*^ = 0); shading: bistable coexistence window. Hysteresis in ascending versus descending frequency ramps is a testable prediction. **(B)** Inhibitory steady-state rate for *f* ≥ *f*_*b*2_ ≈ 140 Hz (note the silent epoch [*f*_th_, *f*_*b*2_]). Red squares: simulation; black dashed: analytical prediction Equation (2). **(C)** Time trace at *f* = 160 Hz *> f*_th_: excitatory suppression and elevated inhibitory activity, consistent with the analytical prediction. **(D)** Phase diagram in the (*f, β*) plane for the default coupling parameters, with boundary lines *f*_th_(*β*) (solid) and *f*_low_(*β*) (dashed) shown for three values of the effective I→E coupling 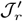 (see legend). Red region: only the high-activity state exists; DBS cannot quench E firing. Yellow region: bistable — both states are stable and the outcome depends on history. Blue region: only the silent E state exists — this is the therapeutic window. White dashed/dotted lines mark the clinical stimulation range 90 Hz/185 Hz. Black circle: *f*_th_ at the default parameter values. Blue squares: the nine DBS frequencies tested in Fig. 3, plotted at the fitted depression strength 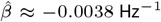. Green triangle: *f*_osc_ = 50 Hz at *β* = −0.003 Hz^*−*1^, the depression strength used in Fig. 4; the dotted green line spans the full frequency range swept in that figure. Both operating points fall well inside the active region at this panel’s default coupling, far from *f*_th_. Parameters: *τ* = *τ*^*′*^ = 0.2 s, *J*_0_ = 0.3, *J*_r_ = 0.4, 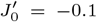, *η* = 0.1, Δ*t* = 0.1 ms; *f*_low_ ≈ 11 Hz, *f*_th_ ≈ 104 Hz, *f*_*b*2_ ≈ 140 Hz.

The coexistence of the high- and low-activity branches over the interval [*f*_low_, *f*_th_] predicts stimulus hysteresis: ascending and descending frequency ramps should yield different transition points (Fig. 2A). This prediction is testable with frequency-ramp intraoperative protocols and provides a behavioral signature that would distinguish the BEB mechanism from purely monostable accounts. Figure 2D maps these three dynamical regions — active only (red), bistable (yellow), and E-suppressed (blue) — in the (*f, β*) parameter plane, for three values of the effective inhibitory feedback coupling 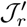. The blue region is the therapeutic target: only here does stimulation guarantee quenching of E activity regardless of initial state. Stronger depression (more negative *β*) or weaker inhibitory coupling shifts *f*_th_ downward and widens the therapeutic window, whereas insufficient depression (| *β* | too small) places *f*_th_ above the practical stimulation range. For context, the operating points used elsewhere in this study are overlaid on the same diagram, evaluated with this panel’s representative default coupling rather than each nucleus’s own fitted parameters: the nine VIM stimulation frequencies and the depression strength fitted in Fig. 3, and the (*β, f*_osc_) condition examined in Fig. 4, both sit well inside the active region, far from *f*_th_ at these *β* values — illustrating, independently of the nucleus-specific coupling fit, why such weak depression generically places the BEB beyond the clinically tested range.

**Fig. 3.**
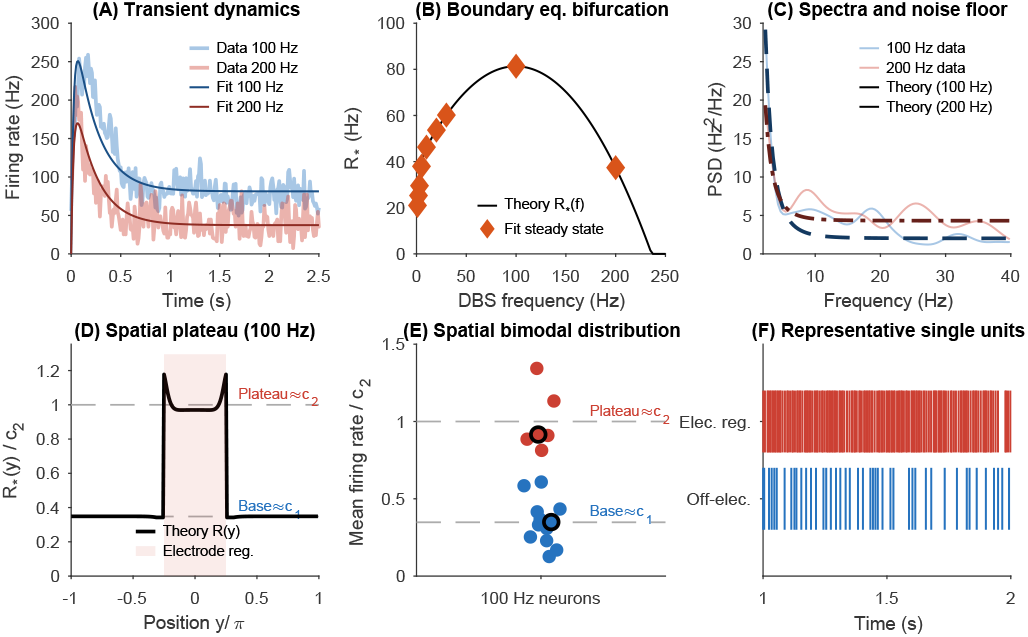
Model comparison with intraoperative VIM recordings. **(A)** Population-averaged instantaneous firing rate (PSTH, 10 ms bins) at 100 Hz (blue, *n* = 18) and 200 Hz (red, *n* = 10); thick curves: jointly fitted model trajectories (fit simultaneously to all nine DBS frequencies 1–200 Hz). **(B)** Steady-state BEB curve: theoretical *R*_*\**_ ((*f*) from fitted parameters (black line); orange diamonds: empirical time-averaged rates at all nine frequencies (*t >* 2 s window). *R*_*\**_ ((*f*) rises then turns over within the tested range, the same downward-concave rectified quadratic identified in Fig. 1; the BEB threshold itself lies beyond 200 Hz, consistent with VIM as a gain-modulation nucleus. **(C)** Welch PSD (2–40 Hz); solid: empirical (100 Hz blue, 200 Hz red); dashed/dotted: LNA prediction from PSD expression given in the SI Appendix with overdamped denominator (Δ > 0, real eigenvalues), scaled to empirical peak above estimated Poisson noise floor. The monotone theoretical shape is a genuine first-principles prediction: *α <* 0 (LNA valid) and the absence of a resonant peak is consistent with VIM not exhibiting autonomous beta oscillations (see text). **(D)** Theoretical spatial profile *R*_*\**_ (*y*) at 100 Hz; shaded: electrode footprint |*y*| ≤ *π/*4; grey dashed lines: plateau level *c*_2_ and baseline level *c*_1_ matched to empirical cluster centroids. **(E)** Distribution of time-averaged firing rates across *n* = 18 neurons at 100 Hz; neurons partitioned by *k*-means into electrode-region (*c*_2_, red) and off-electrode (*c*_1_, blue) groups. Formal bimodality not significant at *n* = 18. **(F)** Spike rasters (*t* ∈ [1, 2] s) for one representative electrode-region neuron (upper) and one off-electrode neuron (lower). *Fitted parameters (9-frequency joint fit, optimized params all*.*mat):* 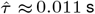, 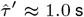 (at optimizer bound; interpreted as an effective population-level inhibitory integration time constant, not a single-cell membrane time constant; see text),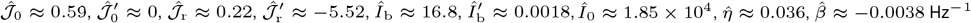

**Fig. 4.**
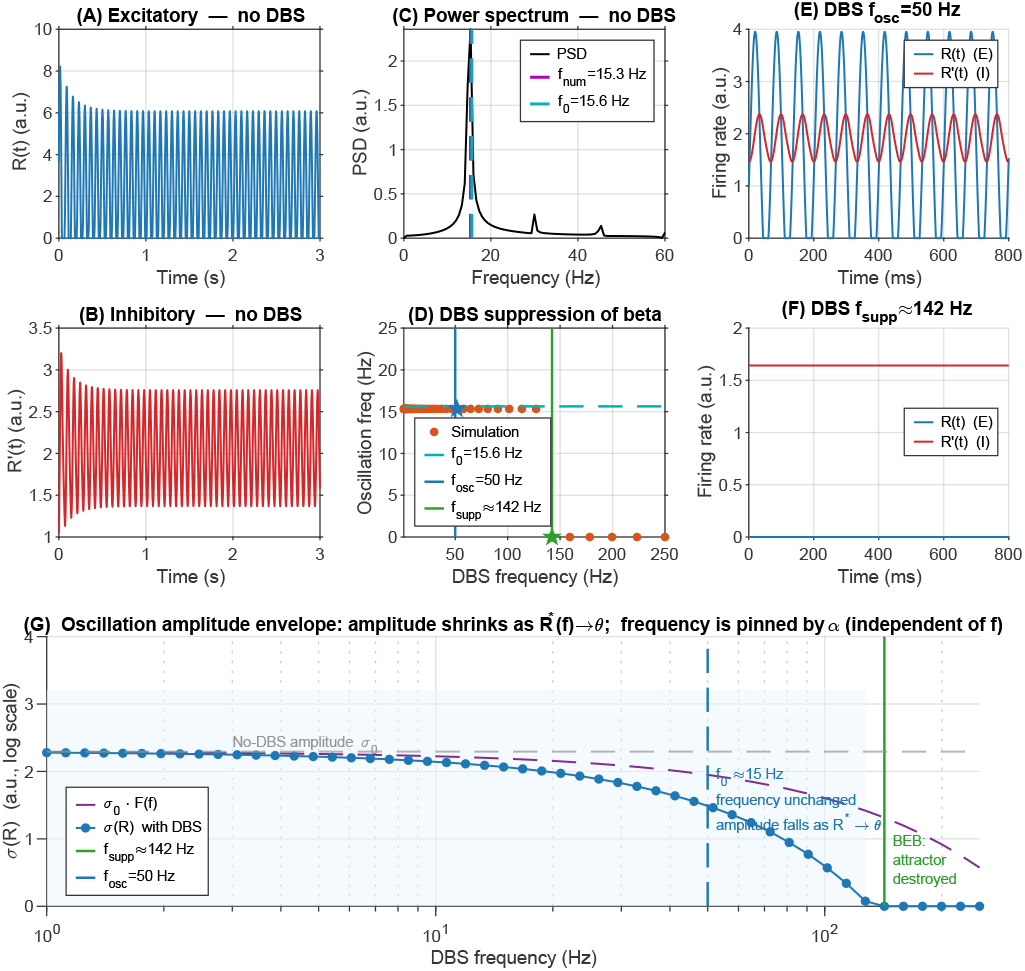
Beta oscillations and DBS suppression via boundary equilibrium bifurcation. **(A)** Excitatory firing rate *R*(*t*) without DBS: sustained limit-cycle oscillation in the beta band. **(B)** Inhibitory rate *R*^*′*^(*t*) without DBS; lags excitatory by ∼ *π/*2, consistent with the E→I→E feedback loop. **(C)** One-sided PSD; magenta dashed: numerically identified peak *f*_num_ ≈ 15.3 Hz; cyan dashed: analytical prediction *f*_0_ ≈ 15.6 Hz (∼2% discrepancy). **(D)** Dominant oscillation frequency vs. DBS frequency. The frequency is constant up to *f*_th_ and drops to zero abruptly, consistent with BEB suppression (not a Hopf shift); *α* (Equation (3)) is independent of *f*. **(E**,**F)** Time traces at *f*_osc_ = 50 Hz (persistent limit cycle) and *f*_supp_ ≈ 142 Hz (fully suppressed), confirming the all-or-nothing character of the transition. **(G)** RMS oscillation amplitude *σ*(*R*) (log scale) versus DBS frequency. The amplitude decays continuously as *f* increases because the limit-cycle orbit shrinks in proportion to *R*_*\**_ (*f*) − *θ* → 0, while the oscillation frequency remains pinned at *f*_0_ (panel D); this decay is entirely geometric — *α* is independent of *f* — not a change in the decay rate of the focus. Purple dashed: *σ*_0_ *F* (*f*), the amplitude expected under the (incorrect) hypothesis that the oscillation simply rescales with the linear drive attenuation; *σ*(*R*) tracks this line at low *f* but falls increasingly below it approaching *f*_th_, confirming that the collapse is governed by proximity to threshold rather than by the magnitude of the drive itself. The grey dashed line marks the no-DBS reference amplitude; the blue shaded region indicates the range of DBS frequencies over which the oscillation is present. Because neural recordings are contaminated by background noise with standard deviation *σ*_noise_, the oscillation becomes undetectable once *σ*(*R*) ≲ *σ*_noise_, well before the formal BEB at *f*_supp_. The observable suppression threshold is therefore set by a signal-to-noise condition on the shrinking orbit, not solely by the bifurcation point itself. Parameters: *τ* = 0.2 s, *τ*^*′*^ = 0.16 s, 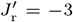, *l* =π/3 *β* = −0.003 Hz^*−*1^. Analytical: *α* ≈ 0.63 s^*−*1^, *f*_0_ ≈ 15.6 Hz.

### Comparison with intraoperative VIM recordings

We next tested whether the same mean-field model, fit to a single nucleus, reproduces the transient and steady-state dynamics observed in intraoperative recordings, rather than only the analytical bifurcation structure derived above. We compared the bi-population ODE model (Equations (7) and (8)) against single-unit recordings from *n* = 18 VIM thalamic neurons at 100 Hz DBS and *n* = 10 neurons at 200 Hz DBS, with 9 neurons recorded at both frequencies (3, 17). The model was fit jointly to all nine stimulation frequencies (1, 2, 3, 5, 10, 20, 30, 100, and 200 Hz) by minimizing a frequency-proportional weighted sum of squared residuals against the population-averaged firing rate trajectory (`fmincon`, 10,000 evaluations; SI Appendix, Section D). Fitting 11 parameters to nine population-averaged time series is underdetermined; these results therefore demonstrate dynamical plausibility and internal consistency rather than constituting a unique parameter identification.

The fitted model trajectories reproduce the transient onset and steady-state level across all nine stimulation frequencies (Fig. 3A). The fitted frequency-attenuation coefficient 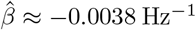 is within the range measured for glutamatergic short-term depression at corticosubthalemic and thalamocortical synapses (18). The steady-state BEB curve from the fitted parameters (Fig. 3B) rises through low and intermediate frequencies and then turns over, declining toward 200 Hz; this rise-then-fall shape is the same rectified-quadratic form established in Fig.1 (downward concavity, 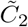, evaluated over a wider frequency range than the 1–100 Hz window used for the cross nuclei classification. A Wilcoxon signed-rank test on the 9 paired neurons confirms a statistically significant rate reduction from 100 to 200 Hz (*p* = 0.0039), consistent with the descending arm of this curve. For VIM — classified here as a gain-modulation nucleus — the analytical BEB threshold *f*_th_ itself falls outside the stimulation range tested (1–200 Hz) under the fitted parameters; the model therefore predicts this smooth, threshold-free rise-then fall modulation in this nucleus, not a sharp threshold suppression.

Welch PSD estimates of the demeaned population PSTH show a spectral elevation in the beta band under both conditions; the 200 Hz condition has lower overall spectral amplitude than the 100 Hz condition (Fig.3C) Three points govern the analytical comparison. First, the LNA requires only *α <* 0 (stable fixed point), a condition that *is satisfied* at the fitted parameters; the exact PSD formula therefore applies. Second, the fitted discriminant Δ > 0: the system is overdamped, with real negative eigenvalues *λ*_1,2_ = *α ± κ*. The PSD denominator evaluates to 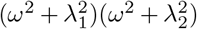, giving a monotone, peak-less spectrum. This is *consistent* with VIM thalamus not generating autonomous beta oscillations; the spectral elevation in the data reflects input-driven activity, not a network resonance. The spectral overlay in panel C is derived directly from the fitted eigenvalues — the shape is fixed once *τ* , *τ* ^′^, and the coupling parameters are known — and is scaled only to match the empirical peak amplitude above an estimated noise floor. It is therefore a parameter-free prediction of the spectral shape, not a separate fit to the PSD data. Third, the spectral sharpness index *C* requires Δ < 0 (a resonant frequency) and is not applicable here; no claim is made about Hopf proximity from the VIM data.

The large fitted inhibitory time constant 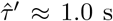(at its optimizer upper bound) reflects the underdetermination of the inhibitory dynamics from excitatory rate data alone. Rather than indicating an implausibly slow single-cell time constant, this value should be interpreted as an effective population-level integration time that encapsulates the multi-synaptic feedback loop through which thalamic inhibition operates; population-level effective time constants are routinely larger than cellular ones (19).

Neuron-to-neuron variability in time-averaged firing rates at 100 Hz (Fig. 3E) is qualitatively consistent with the bimodal spatial profile predicted by the model — neurons near the electrode should fire at a higher mean rate than those outside the stimulated zone. Disentangling the contributions of electrode proximity and local inhibitory feedback requires simultaneous distance measurements and feedback estimates, which are not available in the present dataset. Formal bimodality was not significant at *n* = 18.

### Endogenous beta oscillations and DBS-induced suppression

The same E–I circuit that produces frequency-selective rate suppression also supports endogenous oscillations, connecting the BEB mechanism directly to the pathological beta synchrony that DBS is intended to attenuate. In the high-activity regime, linear stability analysis around the active fixed point yields eigenvalues *λ*_1,2_ = *α* ± *iω*_0_ with:

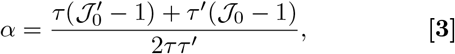

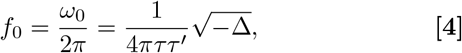

where the discriminant is

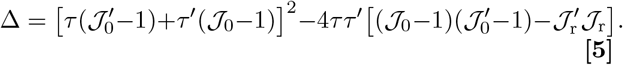

Oscillatory dynamics require Δ < 0. When *α >* 0 (unstable focus), the ReLU nonlinearity bounds the growing oscillation and a stable limit cycle forms in the beta band (Fig. 4A,B). The analytical prediction *f*_0_ ≈15.6 Hz matches the numerically identified peak at *f*_num_ ≈15.3 Hz (~2% discrepancy, consistent with linearizing around an unstable focus rather than the true limit cycle; Fig. 4C).

The real part *α* is determined solely by coupling strengths and synaptic time constants (Equation (3)) entirely independent of stimulation frequency: *F*(*f)* multiplies only the external drive, leaving the recurrent coupling matrix and therefore the Jacobian of the linearized dynamics unchanged at all stimulation frequencies. The Hopf boundary condition *α* = 0 therefore cannot crossed by varying *f*. DBS suppresses beta oscillations not by shifting the Hopf boundary but by eliminating the active fixed point that sustains the limit cycle. As *f* increases past *f*_th_, the active fixed point reaches the ReLU boundary and ceases to exist via the BEB, removing the oscillatory attractor entirely (Fig. 4D–F). Approaching this point, the limit-cycle orbit shrinks continuously as *R*_*\**_ (*f*) −θ *→*0, reducing the oscillation amplitude *σ(R)* while leaving the frequency pinned at *f*_0_; once *σ*(*R*) falls below the background noise floor, the oscillation becomes undetectable in recordings even though the attractor has not yet been formally destroyed (Fig. 4G). This geometric origin is made explicit by comparing *σ*(*R*) against the amplitude expected if the oscillation simply tracked the linear attenuation of the drive itself, *σ*_0_ *F* (*f*): the two coincide at low *f* , but *σ*(*R*) falls increasingly below *σ*_0_ *F*(*f)*as *f → f*_th_ and collapses to zero discontinuously at *f*_th_ (denoted *f*_supp_ ≈142 Hz for this parameter set, to flag that it is the oscillation-suppression point rather than the general BEB threshold reported in Fig. 2), while *σ*_0_ *F* (*f*) remains well above zero at that same frequency. The suppression is therefore not a simple rescaling of the input but a genuinely nonlinear consequence of the fixed point approaching threshold. The predicted signature — oscillation frequency pinned near *f*_0_ up to *f*_supp_, then dropping discontinuously to zero — is qualitatively distinct from gradual Hopf-shift suppression and is directly testable in single DBS-ramp experiments.

In the sub-critical regime (*α <* 0, stable focus), stochastic fluctuations drive resonant activity near ω_0_ The linear noise approximation yields a Lorentzian PSD with FWHM bandwidth Δ*f*_FWHM_ = |*α*| */π*, giving the spectral sharpness index (SI Appendix, Section C.4):

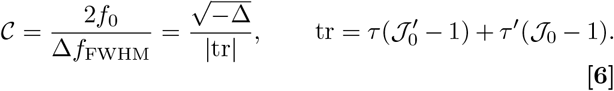

*C* diverges as the Hopf boundary is approached (tr → 0−) and is computable from any LFP recording that resolves both the peak frequency and the bandwidth, without requiring a model fit. It therefore provides a direct, parameter-free readout of how close the circuit is to the oscillatory instability.

### Spectral criticality and spatial DBS footprint

Progressive narrowing of the beta LFP peak is predicted as *J*_0_ approaches the Hopf boundary 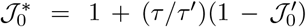from below. The bandwidth Δ*f*_FWHM_ = |*α*|*/π* decreases as 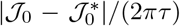 near 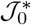 (SI Appendix, Section C.5). Beta-band synchronization in the STN is well established to correlate with parkinsonian motor impairment (20); the present model offers one candidate mechanistic account of this relationship, in which disease progression shifts the circuit toward the Hopf boundary and thereby narrows and sharpens the beta peak, though the specific link between dopaminergic degeneration and changes in effective recurrent coupling remains to be established empirically. Figure 5A–C confirms this across 35 parameter combinations: numerically measured FWHM values track *α /π* with negligible systematic error, validating the linear noise approximation in the sub-critical regime and establishing that is measurable from LFP recordings without fitting a network model. The predicted narrowing rate *d*(Δ*f*_FWHM_)*/d* _0_ = 1*/*(2*πτ*) requires an independent estimate of *τ* from slice electrophysiology to become a quantitative biomarker of disease progression.

**Fig. 5.**
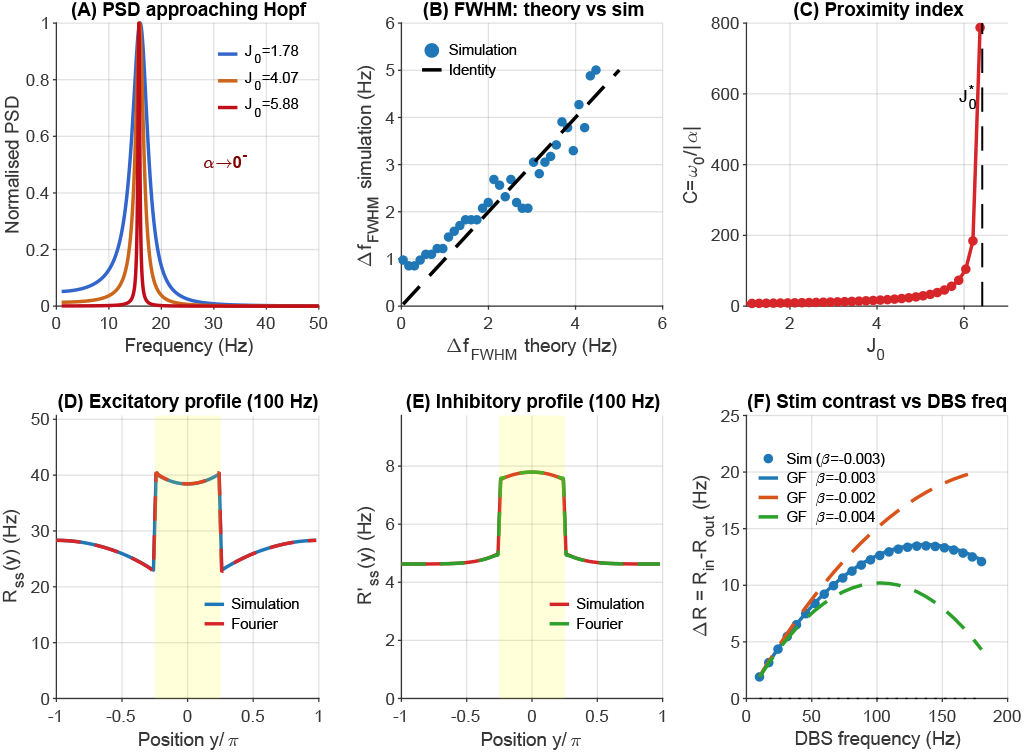
Spectral criticality and spatial DBS footprint. **(A)** Normalised analytical PSD at three values of *J*_0_ approaching the Hopf boundary from below (sub-critical, *α <* 0 throughout). Spectral peak sharpens and narrows as *α* → 0^*−*^, with bandwidth Δ*f*_FWHM_ = |*α*|*/π*. **(B)** FWHM from 35 stochastic simulations (*T* = 60 s Welch) versus analytical prediction |*α*|*/π*; dashed: identity. Quantitative agreement validates the linear noise approximation across the sub-critical regime. **(C)** Spectral sharpness index C = *ω*_0_ */*|*α*| diverges at 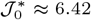 (dashed vertical), providing a parameter-free biomarker of Hopf proximity computable directly from LFP data. **(D**,**E)** Excitatory and inhibitory steady-state spatial profiles at *f* = 100 Hz, electrode footprint *d*_stim_ = *π/*2 (yellow shading). Blue/red: full network simulation; dashed: Fourier decomposition (500 modes). The two profiles have distinct spatial decay constants *κ*_1,2_ set by eigenvalues of the coupling matrix. **(F)** Stimulus contrast Δ*R* = ⟨*R*_ss_⟩in − ⟨*R*_ss_⟩_out_ versus DBS frequency, for the default depression strength *β* = −0.003 (blue) and two bracketing values *β* = −0.002 and *β* = −0.004 (orange/green dashed, Green’s-function theory only). Contrast rises to a maximum at an intermediate frequency *f*_peak_ and then declines; weaker depression shifts *f*_peak_ to higher frequencies and increases the peak contrast. Blue circles: network simulation (default *β*); solid/dashed lines: Fourier/Green’s-function theory. All panels: *l* = *π/*3, *d*_stim_ = *π/*2, *β* = −0.003 Hz^*−*1^ (default), *f* = 100 Hz for panels D,E.

Under focal stimulation (electrode footprint *d <* 2*π*), the periodic Green’s function solution (SI Appendix, Section B.5) gives the exact steady-state spatial profile *R*_ss_(*y*), imposing continuity of the spatial derivative and a step-jump in synaptic input at the electrode boundary. Both the Fourier decomposition (500 modes) and the Green’s function solution reproduce the full *N* = 100 network simulation for decay lengths spanning an order of magnitude (Fig. 5D,E; SI Appendix, Fig. S2). Stimulus contrast Δ*R* = ⟨*R*_ss_ ⟩_in_ − ⟨*R*_ss_ ⟩ _out_ rises with DBS frequency, reaches a maximum at an intermediate frequency *f*_peak_, and then declines over the rest of the plotted range (Fig. 5F) as inhibitory activity increasingly spreads beyond the electrode footprint; for sufficiently strong depression this decline continues to a sign reversal at higher frequencies than shown here — a generic consequence of center-surround connectivity, in which direct excitatory drive falls off less steeply with distance than recurrent inhibitory drive, eventually producing a ring of net suppression flanking the directly activated region. The spatial decay constants 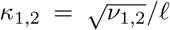(eigenvalues of the coupling matrix divided by the connectivity length) determine the therapeutic activation volume and are accessible from electrode-distance firing-rate profiles.

## Discussion

The analysis supports three related conclusions. The boundary equilibrium bifurcation at *f*_th_ arises from short-term synaptic depression acting through a threshold-linear nonlinearity, and it accounts jointly for the sharp therapeutic threshold and the concurrent rise in GABAergic output with stimulation rate — an empirical pairing that prior models have not derived from a single mechanism. The Hopf instability, by contrast, is governed solely by coupling strengths and synaptic time constants and is invariant under changes in stimulation frequency; DBS therefore suppresses beta oscillations by eliminating the fixed point that sustains the limit cycle, not by shifting the Hopf boundary. Finally, the rectified-quadratic frequency–response is not an empirical fitting form but a structural consequence of *I*_DBS_(*f*) ∝ *f* (1 + *βf*) entering a linear mean-field equation, so the cross nuclei universality inherits its quantitative constraints from the circuit rather than from a phenomenological assumption.

### 0.1 Mechanism and separability

The BEB threshold *f*_th_ is determined by stimulation parameters (*β, I*_0_, Δ*t*) and circuit parameters 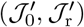 together with thresholds and background drives 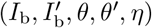 through the quadratic in Equation (1). Because *R ⟶* 0 at the BEB, the *J*_0_*R* and *J*_r_*R* terms (excitatory self-coupling and E→I coupling) vanish identically in the threshold condition, so _0_ and _r_ are absent from Equation (1); 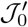 and 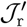 survive because they multiply *R*^′^, which remains nonzero up to threshold. The Hopf boundary 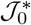 , by contrast, depends exclusively on coupling strengths (including *J*_0_) and membrane time constants and is invariant to changes in stimulation frequency. This algebraic separability — BEB determined only by inhibitory couplings and drive terms, Hopf determined by the full coupling matrix — is a direct consequence of the piecewise-linear ReLU nonlinearity and does not hold in sigmoid-transfer models, where effective gain is state-dependent and hence frequency-modulated. Because *J*_0_ does not enter *f*_th_, pathological PD progression (increasing *J*_0_) and DBS therapy (increasing *f*) operate on orthogonal parameter axes within the same model. This separability offers one account of why DBS can be therapeutic without curing the underlying network pathology: within this model, suppression of pathological firing proceeds through the BEB while the Hopf instability itself is left intact.

#### 02. Relationship to prior models

The conductance-based STN–GPe model of Rubin and Terman (8) provides the most detailed single-cell account of high-frequency DBS effects on pathological bursting; the mean-field framework developed here is complementary, trading single-cell realism for analytical access to population-level bifurcation structure, cross nuclei scaling, and spatial footprint. Biophysical models that incorporate STN projections to the striatum recover beta suppression mechanisms at the network level (21), but do not produce closed-form frequency thresholds or cross nuclei universality relations. Wilson–Cowan E–I models (9, 10) support Hopf bifurcations and beta oscillations, but sigmoid-gain functions preclude BEBs, so they produce smooth rather than threshold frequency responses; abrupt suppression at *f*_th_ is therefore a diagnostic signature of the threshold-linear nonlinearity. Coordinated reset models (11) address patterned desynchronization rather than tonic frequency selectivity. The Lorentzian PSD derived here (exponent 2 far from resonance) is steeper than the 1*/f* -like background typically seen in basal ganglia LFP recordings (22); extending to random sparse connectivity (12, 13) is expected to generate intermediate power-law slopes, since the BEB depends on the mean effective coupling rather than higher-order connectivity moments. The multiplexed coding perspective on synchronized spiking (23) suggests that the two-mode (electrode-region vs. off-electrode) spatial profile predicted by the model may carry distinct information streams, a possibility directly testable with multi-site recordings.

#### 0.3 Cross nuclei organization

The collapsed *R*2 ≥ 0.988 across STN and SNr, and ≥ 0.996 across VIM and RT, is achieved after rescaling by only two empirical parameters per pair: the BEB threshold ratio and the baseline rate ratio. The two-class division itself has a mechanistic interpretation. STN and SNr are classified as threshold-terminated because their fitted BEB threshold *f*_th_ falls within the tested stimulation range (1–100 Hz), where the combination of depression coefficient *β*, pulse amplitude *I*_0_, and background drive *I*_b_ allows the active fixed point to reach the ReLU boundary at clinically accessible frequencies. VIM and RT are gain-modulation nuclei because their fitted parameters place *f*_th_ above the range tested; DBS modulates their output monotonically without reaching the BEB. Whether this reflects genuinely different biophysics (e.g., less short-term depression in thalamocortical vs. basal ganglia synapses) or simply a different operating point within the same model cannot be determined from the present data alone. Two caveats apply. First, a two-parameter rescaling can align any pair of similarly shaped monotone curves; the *R*^2^ value alone does not establish shared underlying computation. Structural content is added by the theory, which constrains the functional form to a rectified quadratic with the specific *f* and *f* ^2^ dependence derived from synaptic depression — a form that is falsifiable and not recoverable by arbitrary two-parameter rescaling of an unconstrained curve. Second, the two-class classification rests on a single intraoperative dataset (3); single-neuron recordings confirm frequency-dependent modulation in these nuclei (17), but whether the two-topology picture holds across disease stages, species, and electrode placements requires independent replication. Neural heterogeneity within each nucleus — known to shape population-level dynamics (7) — may shift the effective BEB threshold and should be incorporated in future extensions.

#### 0.4 Spectral sharpness as a biomarker

The closed-form expression for *C* (Equation (6)) provides a candidate mechanistic account of the beta-band synchronization strength known to correlate with parkinsonian motor impairment (20), via the narrowing and sharpening of the besharpness index *C* peak predicted as the circuit approaches the Hopf boundary. The spectral sharpness index plays the role of a quality factor: it diverges at the Hopf boundary as *α →* 0−, mirroring the critical slowing-down identified in stochastic threshold-crossing problems (24). The predicted bandwidth narrowing rate *d*(Δ*f*_FWHM_)*/dJ* _0_ = − 1*/*(2*πτ*) requires an independent estimate of *τ* from slice electrophysiology to serve as a quantitative disease-progression biomarker. *C* applies only in the sub-critical regime (*α <* 0); it diverges at the Hopf boundary and must not be applied to data recorded during frank limit-cycle oscillations, nor conflated with the 1*/f* exponent of self-organized criticality. The Lorentzian PSD derived here predicts 1*/f* ^2^ tails far from resonance, which is steeper than the broad 1*/f* -like spectral backgrounds observed in basal ganglia LFP recordings (22). This discrepancy is expected: Lorentzian spectra arise from mean-field kinetics with a single pair of complex eigenvalues, and extending to random connectivity generates a distribution of eigenvalues that produces shallower power-law behavior (12, 13).

#### 0.5 Limitations

The linear frequency-attenuation factor *F* (*f*) = 1 + *βf* , which approximates the steady-state limit of the Tsodyks–Markram model (18, 25), neglects pulse-by-pulse vesicle recovery kinetics. Under dynamic depression, the effective drive at *f*_th_ depends on the vesicle recovery time constant *τ*_rec_, and whether the sharp BEB structure survives this substitution requires further analysis; it is plausible, because the BEB depends on the zero crossing of the steady-state fixed point, not on transient dynamics, but this has not been verified. The VIM analysis is necessarily exploratory given sample size: fitting 11 parameters jointly to population-averaged PSTHs from 18 neurons at nine frequencies is underdetermined, and the effective inhibitory time constant 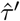 is not well resolved from rate-coded excitatory output alone. As noted above, the large fitted 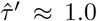 s is best read as this effective population-level time, not a single-cell constant. Extending to simultaneous multi-unit recordings with known electrode-to-neuron distances would substantially improve identifiability. The 1D ring geometry is adequate for the frequency-selectivity and universality results, but quantitative dosimetry — predicting the spatial extent of therapeutic suppression for a given electrode geometry and amplitude — requires three-dimensional extensions. Finally, coupling strengths are treated as fixed constants without explicit links to dopamine concentration or disease state; the connection between *J*_0_ and PD severity is therefore interpretive, and mechanistic predictions about disease progression would require pharmacological modulation of synaptic gains in controlled experiments.

#### 0.6 Falsifiable predictions

Seven predictions distinguish the BEB mechanism from alternatives and are testable with existing or near-term experimental tools. *Hysteresis*: ascending and descending DBS frequency ramps should produce distinct transition points, with the descending transition occurring at *f*_low_, below the ascending transition at *f*_th_, reflecting bistability in [*f*_low_, *f*_th_]. *Inhibitory slope discontinuity*: the inhibitory firing rate should be silent in [*f*_th_, *f*_*b*2_] and then rise linearly above *f*_*b*2_, not from *f*_th_ itself. *Oscillation pinning*: beta frequency should remain near *f*_0_ up to *f*_supp_ and drop to zero discontinuously there, not drift toward zero. *Beta narrowing*: LFP beta bandwidth should decrease linearly with disease severity at rate 1*/*(2*πτ*) per unit increase in recurrent coupling, verifiable by longitudinal LFP recordings paired with estimates of *τ* from in-vitro physiology. *Bimodal spatial firing*: focal DBS should produce a bimodal firing-rate distribution across recorded neurons, with the two modes tracking electrode proximity, and the mode separation determined by the connectivity eigenvalues *κ*_1,2_. *Cross nuclei parameter-free prediction*: once the full frequency–response of one nucleus in a circuit pair is characterized, the partner’s response is predicted without additional parameters via the rescaling relations in SI Appendix, Section A.4. *Footprint scaling*: the therapeutic activation radius should scale with the connectivity decay length *l*, measurable from anatomical tract-tracing or electrode-distance firing-rate profiles, providing a route to patient-specific dosimetry.

Each prediction is testable within existing intraoperative or in-vitro paradigms and produces a clear binary outcome that can discriminate the BEB account from monostable and Hopf-shift alternatives.

## Materials and Methods

### Model equations

The E–I CANN comprises two populations (*N* = 100 neurons each) on a periodic ring *y ∈* [ −*π, π*). The excitatory–inhibitory labelling is adopted for anatomical concreteness; it is a notational convenience rather than a constraint on the coupling signs. Each of the four effective weights 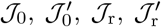may take any sign in data-driven fits, and a negative *J*_0_ or positive 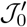 are both admissible. Default parameters are listed in SI Appendix, Table S1. Synaptic inputs obey:

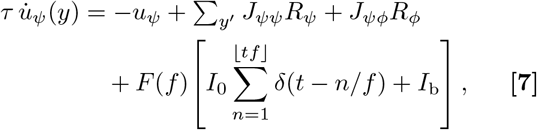

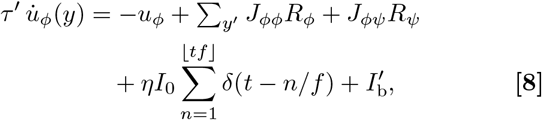

with *R*_*ψ*_ = max(*u*_*ψ*_ −*θ*, 0), *R*_*ϕ*_ = max(*u*_*ϕ*_ −*θ*^′^, 0), *F* (*f*) = max(1 + *βf*, 0), and *J*_*uv*_ (−*y*, −*y*^′^) = *J*_*uv*_ exp(− |*y* −*y*^′^ |_per_*/l*). The transfer function is a threshold-linear (ReLU) approximation to a saturating Michaelis–Menten response, valid when inputs remain well below the saturation rate *R*_max_ (SI Appendix, Section E). Delta pulses are represented numerically as rectangular pulses of amplitude *I*_0_ and duration Δ*t* = 10−4 s. The factor *F* (*f*) appears on the excitatory drive only; the inhibitory drive is not subject to the same degree of frequency-dependent attenuation because GABAergic vesicle pools are less depleted at high stimulation rates (16, 26). The linear approximation F (f) ≈ 1 − Uτ_rec_f is the first-order Taylor expansion of the Tsodyks–Markram steady-state efficacy (1+f τ_rec_U)−1, with β ≡ −Uτ_rec_ (18, 25).

### Analytical derivations

Uniform steady states, bistability conditions, and the BEB threshold (Equation (1)) are derived in SI Appendix, Section A. Closed-form expressions for *f*_0_ and (Equations (4) and (6)) are derived in SI Appendix, Section C. The periodic Green’s function spatial solution and the Fourier mode decomposition (SI Appendix, Section B) are validated against *N* = 100 network simulations in SI Appendix, Figs. S1–S2. Nucleus-specific model instantiations (VIM/RT, STN/GPe, SNr) and the inhibitory-halo spatial effect are detailed in SI Appendix, Sections F–G.

### Numerical simulations

Forward Euler integration (Δ*t*_sim_ = 10−3 s) was used for deterministic simulations. Stochastic simulations (PSD analysis) used Δ*t* = 10−4 s, *T* = 60 s, and additive Gaussian noise (*σ* = 0.3 a.u.) applied independently to each population after each Euler step. Spectral estimation used Welch’s method (Hann window, *L* = 2048, overlap 1024, *N*_fft_ = 8192, *f*_*s*_ = 1 kHz after downsampling by a factor of 10 from the simulation rate).

### Data and fitting

Multi-nucleus frequency–response data are from Milosevic et al. (3), who recorded single-unit activity during intraoperative DBS across STN, SNr, VIM, and RT in patients undergoing electrode implantation surgery, under institutional ethics approval obtained for the original recordings. The raw single-unit VIM-DBS recordings used in Fig. 1 are publicly available (3); the processed dataset used for the transient and steady-state dynamics fit in Fig. 3 was obtained from Paraskevopoulos et al. (17). Model fitting used `fmincon` (MATLAB R2023b, SQP algorithm, 10,000 function evaluations, single warm start from default parameters) minimizing a frequency-proportional weighted sum of squared residuals *w*_*k*_ = *f*_*k*_*/f*_min_ between model output and population-averaged PSTH across all nine stimulation frequencies (1–200 Hz). The weighting scheme prioritizes agreement at clinically relevant high frequencies without discarding information from low-frequency conditions.

## Code availability

Analysis code and the fitted parameter file (`optimized params all.mat`;) are available at https://github.com/xma-samuel/dbs-cann.

## ACKNOWLEDGMENTS

We thank Zoe Paraskevopoulos for her assistance with processing the intraoperative VIM single-unit recording data. We also thank members of the Lankarany laboratory for helpful discussions.

## References

1. CC McIntyre, M Savasta, L Kerkerian-Le Goff, JL Vitek, Uncovering the mechanism(s) of action of deep brain stimulation: activation, inhibition, or disruption? Clin. Neurophysiol. 115, 1239–1248 (2004).

2. WJ Neumann, LA Steiner, L Milosevic, Neurophysiological mechanisms of deep brain stimulation across spatiotemporal resolutions. Brain 146, 4456–4468 (2023).

3. L Milosevic, et al., A theoretical framework for the site-specific and frequency-dependent neuronal effects of deep brain stimulation. Brain Stimul. 14, 807–821 (2021).

4. JS Brittain, P Brown, Oscillations and the basal ganglia: Motor control and beyond. NeuroImage 85, 637–647 (2014).

5. P Brown, Oscillatory nature of human basal ganglia activity: relationship to the pathophysiology of Parkinson’s disease. Mov. Disord. 18, 357–363 (2003).

6. J Li, et al., Differential synaptic depression mediates the therapeutic effect of deep brain stimulation. Nat. Neurosci. 28, 2575–2587 (2025).

7. A Hutt, S Rich, TA Valiante, J Lefebvre, Intrinsic neural diversity quenches the dynamic volatility of neural networks. Proc. Natl. Acad. Sci. USA 120, e2218841120 (2023).

8. JE Rubin, D Terman, High frequency stimulation of the subthalamic nucleus eliminates pathological thalamic rhythmicity in a computational model. J. Comput. Neurosci. 16, 211–235 (2004).

9. HR Wilson, JD Cowan, Excitatory and inhibitory interactions in localized populations of model neurons. Biophys. J. 12, 1–24 (1972).

10. S Coombes, Á Byrne, Next generation neural mass models in Nonlinear Dynamics in Computational Neuroscience. (Springer), pp. 1–16 (2019).

11. PA Tass, A model of desynchronizing deep brain stimulation with a demand-controlled coordinated reset of neural subpopulations. Biol. Cybern. 89, 81–88 (2003).

12. K Rajan, LF Abbott, Eigenvalue spectra of random matrices for neural networks. Phys. Rev. Lett. 97, 188104 (2006).

13. S Ostojic, Two types of asynchronous activity in networks of excitatory and inhibitory spiking neurons. Nat. Neurosci. 17, 594–600 (2014).

14. Si Amari, Dynamics of pattern formation in lateral-inhibition type neural fields. Biol. Cybern. 27, 77–87 (1977).

15. S Wu, KYM Wong, CCA Fung, Y Mi, W Zhang, Continuous attractor neural networks: Candidate of a canonical model for neural information representation. F1000Research 5, 156 (2016).

16. Z Xu, et al., Deep brain stimulation alleviates parkinsonian motor deficits through desynchronizing GABA release in mice. Nat. Commun. 16, 3726 (2025) Article 3726.

17. Z Paraskevopoulos, et al., Frequency-dependent inhibition during deep brain stimulation of thalamic ventral intermediate nuclei. J. Neurosci. 46, e1859252026 (2026).

18. MV Tsodyks, H Markram, The neural code between neocortical pyramidal neurons depends on neurotransmitter release probability. Proc. Natl. Acad. Sci. 94, 719–723 (1997).

19. P Dayan, LF Abbott, Theoretical Neuroscience: Computational and Mathematical Modeling of Neural Systems. (MIT Press, Cambridge, MA), (2001).

20. AA Kühn, A Kupsch, GH Schneider, P Brown, Reduction in subthalamic 8–35 Hz oscillatory activity correlates with clinical improvement in Parkinson’s disease. Eur. J. Neurosci. 23, 1956–1960 (2006).

21. EM Adam, EN Brown, N Kopell, MM McCarthy, Deep brain stimulation in the subthalamic nucleus for Parkinson’s disease can restore dynamics of striatal networks. Proc. Natl. Acad. Sci. USA 119, e2120808119 (2022).

22. C Wiest, et al., The aperiodic exponent of subthalamic field potentials reflects excitation/inhibition balance in Parkinsonism. eLife 12, e82467 (2023).

23. M Lankarany, D Al-Basha, S Ratté, SA Prescott, Differentially synchronized spiking enables multiplexed neural coding. Proc. Natl. Acad. Sci. USA 116, 10097–10102 (2019).

24. T Taillefumier, MO Magnasco, A phase transition in the first passage of a Brownian process through a fluctuating boundary with implications for neural coding. Proc. Natl. Acad. Sci. USA 110, E1438–E1443 (2013).

25. LF Abbott, JA Varela, K Sen, SB Nelson, Synaptic depression and cortical gain control. Science 275, 220–224 (1997).

26. S Chiken, A Nambu, High-frequency pallidal stimulation disrupts information flow through the pallidum by GABAergic inhibition. J. Neurosci. 33, 2268–2280 (2013).

