## Supplementary Information for "Bifurcation Structure and Cross Nuclei Universality Govern Frequency-Selective Deep Brain Stimulation"

Table S1: **Table S1.** Default model parameters for spatially uniform analysis. Per-figure deviations are stated in each caption. Scaled coupling strengths $\mathcal{J}_0 = 2\rho J_0 l$ etc. are computed in SI Appendix, Section A.1.

| Symbol | Description | Default | Units |
|---|---|---|---|
| $N$ | Neurons per population | 100 | – |
| $\tau$ | Excitatory synaptic time constant | 0.2 | s |
| $\tau'$ | Inhibitory synaptic time constant | 0.2 | s |
| $J_0$ | Excitatory recurrent strength | 0.2 | – |
| $J_0'$ | Inhibitory self-coupling | $-0.1$ | – |
| $J_\mathrm{r}$ | E→I coupling | 0.1 | – |
| $J_\mathrm{r}'$ | I→E coupling | $-3.0$ | – |
| $l$ | Spatial decay length (all kernels) | $\pi/3$ | rad |
| $\theta$ | Excitatory threshold | 10 | a.u. |
| $\theta'$ | Inhibitory threshold | 10 | a.u. |
| $I_\mathrm{b}$ | Excitatory background input | 200 | a.u. |
| $I_\mathrm{b}'$ | Inhibitory background input | 10 | a.u. |
| $I_0$ | DBS pulse amplitude | 5000 | a.u. |
| $\beta$ | Frequency-depression coefficient† | $-0.003$ | $\mathrm{Hz}^{-1}$ |
| $\eta$ | GABAergic DBS fraction | 0.1 | – |
| $\Delta t$ | DBS pulse duration | $10^{-4}$ | s |
| $\sigma_\psi, \sigma_\phi$ | Noise intensities (PSD sims.) | 0.3 | a.u. |

†Default value used for theoretical figures; the VIM joint fit yields $\hat{\beta} \approx -0.0038$ $\mathrm{Hz}^{-1}$ (SI Appendix, Section D.4).

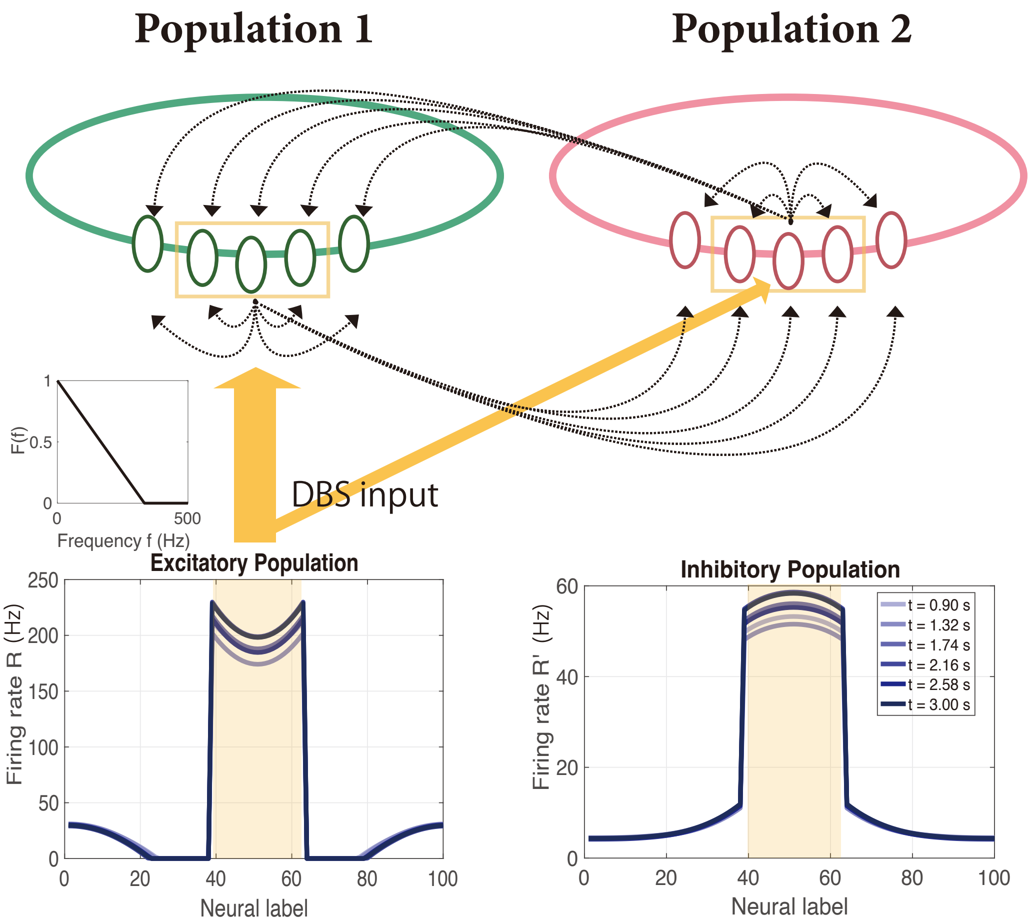


Figure S0: **Fig. S0. Bi-population CANN architecture.** Two neural populations are arranged on one-dimensional circular arrays with periodic boundary conditions. Population 1 (excitatory, green) represents the principal DBS target nucleus (VIM, STN, SNr, or RT); Population 2 (inhibitory, red) represents the interacting GABAergic population within the same circuit loop (RT for VIM thalamus; GPe for STN; collateral GABAergic neurons for SNr). DBS drives Population 1 with frequency-dependent efficacy scaling $F(f) = \max(1 + \beta f, 0)$, $\beta < 0$ (orange arrows; see SI Appendix, Section E for justification); a fraction $\eta$ of the pulse current reaches Population 2 directly. Within each population, recurrent connections decay exponentially with functional distance (black dotted arrows); bidirectional cross-population connections are indicated by curved arrows. *Bottom panels:* representative firing-rate profiles showing a localized activity bump centered at the stimulation site under focal DBS ($d_{\text{stim}} = \pi/2$), computed from the numerical simulation.

### A Spatially Uniform Dynamics: Full Derivation

#### A.1 Spatial Integration of Connection Kernels

We begin by reducing the spatially extended network to an effective point-mass system. Assume that in the spatially uniform regime the synaptic inputs are independent of position: $\psi(y,t) = \psi_0(t)$ and $\phi(y,t) = \phi_0(t)$ for all $y \in [-\pi, \pi)$. The discrete sum over pre-synaptic neurons is replaced by a continuous integral using the neural density $\rho = N/(2\pi)$:

$$\sum_{y'=-\pi}^{\pi} J(y,y')\, R(y') \;\approx\; \rho \int_{-\pi}^{\pi} J_0\, e^{-|y-y'|/l_0}\, R_0\, dy', \tag{A.1}$$

where $R_0 = R(y', t)$ is the spatially uniform firing rate. Factoring out $R_0$ and evaluating the integral:

$$\rho J_0 R_0 \int_{-\pi}^{\pi} e^{-|y-y'|/l_0}\, dy' = 2\rho J_0 l_0 \Big(1 - e^{-\pi/l_0}\Big) R_0, \tag{A.2}$$

where we shifted $u = |y' - y|$ and used the symmetry of the kernel. For the shortest-arc periodic kernel ($|y|_{\text{per}} = \min(|y|, 2\pi - |y|)$, which equals $|y|$ for $|y| \le \pi$), the exact result is

$$\int_{-\pi}^{\pi} e^{-|y-y'|_{\text{per}}/l_0}\, dy' = 2l_0\big(1 - e^{-\pi/l_0}\big). \tag{A.3}$$

For the image-sum (fully periodic) kernel $\sum_{k=-\infty}^{\infty} e^{-|u+2\pi k|/l_0}$, the integral over one period equals exactly $2l_0$, since the individual images tile $(-\infty, \infty)$ without overlap and $\int_{-\infty}^{\infty} e^{-|v|/l_0}\, dv = 2l_0$. In either case, for $l_0 \ll \pi$ both results reduce to $2l_0$, recovering Equation (A.4). For connection decay lengths $l_0 \ll \pi$ (i.e. when most synaptic weight decays within the domain), we have $e^{-\pi/l_0} \approx 0$ and the result simplifies to

$$\rho \int_{-\pi}^{\pi} J_0\, e^{-|y-y'|_{\text{per}}/l_0}\, dy' \;\approx\; 2\rho J_0 l_0 \;\equiv\; \mathcal{J}_0. \tag{A.4}$$

Effective spatial coupling strengths for all four connection types:

$$\mathcal{J}_0 \equiv 2\rho J_0 l_0, \qquad \mathcal{J}'_0 \equiv 2\rho J'_0 l'_0, \tag{A.5}$$

$$\mathcal{J}_\mathrm{r} \equiv 2\rho J_\mathrm{r} l_\mathrm{r}, \qquad \mathcal{J}'_\mathrm{r} \equiv 2\rho J'_\mathrm{r} l'_\mathrm{r}. \tag{A.6}$$

The E–I labelling is a notational convenience; each of these four effective weights may take any sign. In the canonical E–I interpretation, $\mathcal{J}_0 > 0$, $\mathcal{J}'_0 < 0$, $\mathcal{J}_\mathrm{r} > 0$, $\mathcal{J}'_\mathrm{r} < 0$, but data-driven fits may yield atypical signs reflecting the net effect of a multi-synaptic pathway; for example, a disinhibitory circuit can produce an effectively positive $\mathcal{J}'_\mathrm{r}$, and a near-zero $\mathcal{J}'_0$ as found in the VIM fit is fully admissible.

#### A.2 Reduction to Coupled ODEs

Under the uniform ansatz, the governing equations reduce to a pair of first-order scalar ODEs:

$$\tau\,\dot{\psi}_0 = -\psi_0 + \mathcal{J}_0 R_0 + \mathcal{J}'_\mathrm{r} R'_0 + F(f)\big[I_0 f \Delta t + I_\mathrm{b}\big], \tag{A.7}$$

$$\tau'\,\dot{\phi}_0 = -\phi_0 + \mathcal{J}'_0 R'_0 + \mathcal{J}_\mathrm{r} R_0 + \eta I_0 f \Delta t + I'_\mathrm{b}, \tag{A.8}$$

where the periodic pulse train has been replaced by its time-averaged value $f\Delta t$ (valid for $f \gg 1/\tau$), and $R_0 = \max[\psi_0 - \theta, 0]$, $R'_0 = \max[\phi_0 - \theta', 0]$.

#### A.3 Steady-State Solutions and Bistability

At steady state $\dot{\psi}_0 = \dot{\phi}_0 = 0$.

**Low-activity branch ($R_0 = 0,\ R_0' = 0$).**

$$\psi_0^{\text{low}} = F(f)\big[I_0 f \Delta t + I_{\text{b}}\big], \tag{A.9}$$

$$\phi_0^{\text{low}} = \eta I_0 f \Delta t + I_{\text{b}}'. \tag{A.10}$$

This branch is self-consistent whenever $\psi_0^{\text{low}} \le \theta$ and $\phi_0^{\text{low}} \le \theta'$.

**High-activity branch ($R_0 > 0,\ R_0' > 0$).** The fixed-point equations reduce to the $2 \times 2$ linear system $\mathbf{M}\begin{pmatrix}\psi_0 \\ \phi_0\end{pmatrix} = \begin{pmatrix} b_1 \\ b_2 \end{pmatrix}$ with $\mathbf{M} = \begin{pmatrix} 1 - \mathcal{J}_0 & -\mathcal{J}_{\text{r}}' \\ -\mathcal{J}_{\text{r}} & 1 - \mathcal{J}_0' \end{pmatrix}$,

$$b_1(f) = -\mathcal{J}_0\theta - \mathcal{J}_{\text{r}}'\theta' + F(f)\big[I_0 f \Delta t + I_{\text{b}}\big], \tag{A.11}$$

$$b_2(f) = -\mathcal{J}_{\text{r}}\theta - \mathcal{J}_0'\theta' + \eta I_0 f \Delta t + I_{\text{b}}'. \tag{A.12}$$

The system determinant is

$$\Sigma_{\text{sys}} = (1 - \mathcal{J}_0)(1 - \mathcal{J}_0') - \mathcal{J}_{\text{r}}'\mathcal{J}_{\text{r}}. \tag{A.13}$$

Provided $\Sigma_{\text{sys}} \neq 0$, Cramer's rule gives

$$\psi_0^{\text{high}}(f) = \frac{(1 - \mathcal{J}_0')\, b_1(f) + \mathcal{J}_{\text{r}}'\, b_2(f)}{\Sigma_{\text{sys}}}, \tag{A.14}$$

$$\phi_0^{\text{high}}(f) = \frac{\mathcal{J}_{\text{r}}\, b_1(f) + (1 - \mathcal{J}_0)\, b_2(f)}{\Sigma_{\text{sys}}}. \tag{A.15}$$

Substituting $b_1, b_2$ and collecting powers of $f$ yields $R_*(f) \equiv \psi_0^{\text{high}}(f) - \theta = C_0 + C_1 f + C_2 f^2$, with

$$C_0 = \frac{(1 - \mathcal{J}_0')(I_{\text{b}} - \theta) + \mathcal{J}_{\text{r}}'(I_{\text{b}}' - \theta')}{\Sigma_{\text{sys}}}, \tag{A.16}$$

$$C_1 = \frac{\left[(1 - \mathcal{J}_0') + \eta\, \mathcal{J}_{\text{r}}'\right] I_0\, \Delta t + (1 - \mathcal{J}_0')\, I_{\text{b}}\, \beta}{\Sigma_{\text{sys}}}, \tag{A.17}$$

$$C_2 = \frac{(1 - \mathcal{J}_0')\, I_0\, \Delta t\, \beta}{\Sigma_{\text{sys}}}. \tag{A.18}$$

Observe that $C_2 \propto \beta$ and vanishes for $\beta = 0$, consistent with Equation (A.19) below.

#### A.4 Quadratic Frequency–Response: Derivation

For any network in the active linear regime with frequency-dependent synaptic efficacy $F(f) = 1 + \beta f$, the time-averaged DBS input per pulse cycle is

$$I_{\text{DBS}}(f) = F(f)\, I_0\, \Delta t\, f = I_0\, \Delta t\, f + I_0\, \Delta t\, \beta\, f^2, \tag{A.19}$$

which is exactly quadratic in $f$. Because the network dynamics are linear in the drive and temporal averaging is a linear operation, the mean firing rate inherits this form:

$$R_{\text{avg}}(f) = \max\!\big(\tilde{C}_2\, f^2 + \tilde{C}_1\, f + \tilde{C}_0,\ 0\big). \tag{A.20}$$

Therefore, the cross nuclei prediction equations introduced in the main text are

$$R_{\text{STN}}(f) = \mathcal{A} \cdot R_{\text{SNr}}(\mathcal{F}\, f)\,, \quad \mathcal{A} \equiv \frac{C_0^{\text{STN}}}{C_0^{\text{SNr}}}, \quad \mathcal{F} \equiv \frac{f_{\text{th}}^{\text{SNr}}}{f_{\text{th}}^{\text{STN}}} \tag{A.21}$$

and $R_{\text{VIM}}(f) = \mathcal{P}\, R_{\text{RT}}(f) + \mathcal{Q}$, where $\mathcal{P} = \Delta R_{\text{max}}^{\text{VIM}} / \Delta R_{\text{max}}^{\text{RT}}$.

#### A.5 Critical Frequency and Boundary Equilibrium Bifurcation

The critical frequency $f_{\rm th}$ satisfies $\psi_0^{\rm high}(f_{\rm th}) = \theta$. Expanding $b_1(f_{\rm th})$ using $F(f) = 1 + \beta f$ and collecting powers of $f_{\rm th}$ yields the quadratic $Af_{\rm th}^2 + Bf_{\rm th} + C = 0$ with

$$A = (1 - \mathcal{J}_0')\,\beta I_0 \Delta t, \tag{A.22}$$

$$B = (1 - \mathcal{J}_0')(I_0\Delta t + \beta I_{\rm b}) + \mathcal{J}_{\rm r}'\,\eta I_0 \Delta t, \tag{A.23}$$

$$C = (1 - \mathcal{J}_0')(I_{\rm b} - \theta) + \mathcal{J}_{\rm r}'(I_{\rm b}' - \theta'). \tag{A.24}$$

Because $\beta < 0$, $A < 0$. The physically relevant root is

$$f_{\rm th} = \frac{-B - \sqrt{B^2 - 4AC}}{2A}. \tag{A.25}$$

Above $f_{\rm th}$, the inhibitory equation decouples and the inhibitory firing rate grows as

$$R_{\rm ss}'(f) = \frac{\eta I_0 f \Delta t - \theta' + I_{\rm b}'}{1 - \mathcal{J}_0'}, \qquad f \geq f_{b2}, \tag{A.26}$$

where $f_{b2} = (\theta' - I_{\rm b}')/(\eta I_0 \Delta t)$.

#### A.6 Linearization and Hopf Bifurcation Boundary

Perturbing around the high-activity fixed point $(\psi_*, \phi_*)$ gives the Jacobian

$$\mathbf{A} = \begin{pmatrix} (\mathcal{J}_0 - 1)/\tau & \mathcal{J}_{\rm r}'/\tau \\ \mathcal{J}_{\rm r}/\tau' & (\mathcal{J}_0' - 1)/\tau' \end{pmatrix}. \tag{A.27}$$

The eigenvalues are $\lambda_{1,2} = \alpha \pm i\omega_0$ with

$$\alpha = \frac{\rm tr}{2\tau\tau'}, \qquad \omega_0 = \frac{\sqrt{-\Delta}}{2\tau\tau'}, \tag{A.28}$$

where $\mathrm{tr} = \tau(\mathcal{J}_0' - 1) + \tau'(\mathcal{J}_0 - 1)$ and

$$\Delta = \mathrm{tr}^2 - 4\tau\tau'\big[(\mathcal{J}_0 - 1)(\mathcal{J}_0' - 1) - \mathcal{J}_\mathrm{r}'\mathcal{J}_\mathrm{r}\big]. \tag{A.29}$$

A limit cycle arises when $\Delta < 0$ and $\alpha > 0$. The Hopf boundary is defined by $\alpha = 0$:

$$\mathcal{J}_0^* = 1 + \frac{\tau}{\tau'}(1 - \mathcal{J}_0'). \tag{A.30}$$

#### A.7 Transient Response to Pulsatile DBS

Between consecutive pulses at $t_k^+$ and $t_{k+1}^-$:

$$\mathbf{x}(t) = e^{\mathbf{A}(t-t_k)}\mathbf{x}(t_k^+) + \mathbf{A}^{-1}\big[e^{\mathbf{A}(t-t_k)} - \mathbf{I}\big]\mathbf{b}_\mathrm{bg}, \tag{A.31}$$

with instantaneous jump $\mathbf{x}(t_k^+) = \mathbf{x}(t_k^-) + \mathbf{b}_\mathrm{pulse}$ at each pulse. For $\Delta < 0$ (oscillatory regime, complex eigenvalues $\lambda_{1,2} = \alpha \pm i\omega_0$), the matrix exponential is

$$e^{\mathbf{A}t} = e^{\alpha t}\begin{pmatrix} \cos\omega_0 t + \frac{A_{11}-\alpha}{\omega_0}\sin\omega_0 t & \frac{A_{12}}{\omega_0}\sin\omega_0 t \\ \frac{A_{21}}{\omega_0}\sin\omega_0 t & \cos\omega_0 t + \frac{A_{22}-\alpha}{\omega_0}\sin\omega_0 t \end{pmatrix}. \tag{A.32}$$

*Remark* A.1 (Real-eigenvalue case, $\Delta > 0$). When $\Delta > 0$ (overdamped regime), the eigenvalues are real: $\lambda_{1,2} = \alpha \pm \kappa$ with $\kappa = \sqrt{\Delta}/(2\tau\tau') > 0$. The matrix exponential then takes the hyperbolic form

$$e^{\mathbf{A}t} = e^{\alpha t}\begin{pmatrix} \cosh\kappa t + \dfrac{A_{11}-\alpha}{\kappa}\sinh\kappa t & \dfrac{A_{12}}{\kappa}\sinh\kappa t \\ \dfrac{A_{21}}{\kappa}\sinh\kappa t & \cosh\kappa t + \dfrac{A_{22}-\alpha}{\kappa}\sinh\kappa t \end{pmatrix}. \tag{A.33}$$

This form applies whenever $\Delta > 0$, including the VIM fitted parameters (Section D; $\hat{\tau}' = 1.0$ s gives $\Delta > 0$). Both forms are special cases of $e^{\mathbf{A}t} = e^{\alpha t}[f_c(t)\,\mathbf{I} + f_s(t)\,(\mathbf{A} - \alpha\mathbf{I})]$ where

$(f_c, f_s) = (\cos, \sin/\omega_0)$ for $\Delta < 0$ and $(f_c, f_s) = (\cosh, \sinh/\kappa)$ for $\Delta > 0$. The ODE numerical integration used throughout (Euler–Maruyama) does not require this formula explicitly; it is provided for analytical single-pulse response calculations.

### B Spatiotemporal Solution and Steady-State Spatial Profile

#### B.1 Fourier Decomposition of the Integro-Differential Equations

Expanding the spatial fields in cosine series on $[-\pi, \pi)$:

$$\psi(y,t) = \xi_0(t) + \sum_{n=1}^{\infty} \xi_n(t)\cos(ny), \tag{B.1}$$

$$\phi(y,t) = \zeta_0(t) + \sum_{n=1}^{\infty} \zeta_n(t)\cos(ny). \tag{B.2}$$

The exact Fourier coefficient of the periodic exponential kernel is

$$\hat{J}_n = \rho \int_{-\pi}^{\pi} J_0\, e^{-|y|_{\mathrm{per}}/l}\, \cos(ny)\, dy = 2\rho J_0 l\, \frac{1 - (-1)^{|n|} e^{-\pi/l}}{1 + (nl)^2}, \tag{B.3}$$

incurring no wrap-around error. In the limit $l \ll \pi$, this reduces to $\hat{J}_n \approx \mathcal{J}_0/(1 + l^2 n^2)$.

#### B.2 Mode-Dependent System Matrix and Dynamics

The $n$-th mode evolves as $\dot{\mathbf{x}}_n = \mathbf{A}_n \mathbf{x}_n + \mathbf{b}_n(t)$ with

$$\mathbf{A}_n = \begin{pmatrix} \dfrac{1}{\tau}\left(\dfrac{\mathcal{J}_0}{1 + l^2 n^2} - 1\right) & \dfrac{1}{\tau}\dfrac{\mathcal{J}_{\mathrm{r}}'}{1 + l^2 n^2} \\ \dfrac{1}{\tau'}\dfrac{\mathcal{J}_{\mathrm{r}}}{1 + l^2 n^2} & \dfrac{1}{\tau'}\left(\dfrac{\mathcal{J}_0'}{1 + l^2 n^2} - 1\right) \end{pmatrix}. \tag{B.4}$$

Between DBS pulses:

$$\mathbf{x}_n(t) = e^{\mathbf{A}_n(t-t_k)}\mathbf{x}_n(t_k^+) + \mathbf{A}_n^{-1}\left[e^{\mathbf{A}_n(t-t_k)} - \mathbf{I}\right]\mathbf{b}_{n,\mathrm{bg}}, \tag{B.5}$$

with amplitude jumps $\mathbf{x}_n(t_k^+) = \mathbf{x}_n(t_k^-) + \mathbf{b}_{n,\mathrm{pulse}}$ at each pulse.

*Remark* B.1 (Complementary roles of the Fourier and Green's function methods). The Fourier decomposition is optimal for *time-dependent dynamics*: it propagates each spatial mode independently, making it the natural tool for transient responses and spatial propagation of neural oscillations. The Green's function method (Sections B.3–B.5) is optimal for *steady-state analysis*: it yields compact closed-form profiles with algebraic access to spatial decay constants $\kappa_{1,2}$ without mode summation. Both methods achieve excellent quantitative agreement with full-network simulation for all physiologically relevant connectivity decay lengths $l$.

#### B.3 Time-Averaged Steady State and Coupled Spatial ODEs

At high DBS frequencies ($f \gg 1/\tau$), replacing the pulsatile drive with its time-averaged equivalent $\bar{I}^{\mathrm{dbs}}(y) = I_0 f \Delta t\, P(y)$ and setting time derivatives to zero yields the steady-state system. Applying the Green's function operator $\mathcal{L} = (1 - l^2\partial_{yy})$ to both sides reduces the integro-differential equations to

$$l^2 \frac{d^2}{dy^2}\mathbf{u}(y) = \mathbf{M}\mathbf{u}(y) - \mathbf{c}(y), \tag{B.6}$$

where $\mathbf{u}(y) = (\psi_{\mathrm{ss}}(y), \phi_{\mathrm{ss}}(y))^T$, $\mathbf{M} = \begin{pmatrix} 1 - \mathcal{J}_0 & -\mathcal{J}_{\mathrm{r}}' \\ -\mathcal{J}_{\mathrm{r}} & 1 - \mathcal{J}_0' \end{pmatrix}$, and $\mathbf{c}(y)$ is piecewise constant ($\mathbf{c}_{\mathrm{in}}$ inside the electrode, $\mathbf{c}_{\mathrm{out}}$ outside).

#### B.4 Piecewise Solution and Boundary Conditions

The particular solutions on each arc are the spatially uniform fixed points $\mathbf{u}^*_{\text{in}} = \mathbf{M}^{-1}\mathbf{c}_{\text{in}}$ and $\mathbf{u}^*_{\text{out}} = \mathbf{M}^{-1}\mathbf{c}_{\text{out}}$. By symmetry about $y = 0$ (inner arc) and $y = \pm\pi$ (outer arc), the unique admissible piecewise ansatz is:

$$\mathbf{u}(y) = \begin{cases} \mathbf{u}^*_{\text{in}} + \cosh\big(\mathbf{\Lambda} y\big)\, \mathbf{a}, & |y| \le a, \\ \mathbf{u}^*_{\text{out}} + \cosh\big(\mathbf{\Lambda}(\pi - |y|)\big)\, \mathbf{c}, & a < |y| \le \pi, \end{cases} \tag{B.7}$$

where $a = d/2$, $\mathbf{\Lambda} = \mathbf{M}^{1/2}/l$. The boundary conditions at $y = a$ are a jump in $\mathbf{u}$ (reflecting the discontinuous external drive) and continuity of $\partial_y \mathbf{u}$:

$$\cosh\big(\mathbf{\Lambda}_{\text{out}}\big)\mathbf{c} - \cosh\big(\mathbf{\Lambda}_a\big)\mathbf{a} = \mathbf{u}^*_{\text{in}} - \mathbf{u}^*_{\text{out}} - \Delta\mathbf{I}, \tag{B.8}$$

$$\sinh\big(\mathbf{\Lambda}_{\text{out}}\big)\mathbf{c} + \sinh\big(\mathbf{\Lambda}_a\big)\mathbf{a} = \mathbf{0}, \tag{B.9}$$

where $\mathbf{\Lambda}_a = \mathbf{\Lambda} a$, $\mathbf{\Lambda}_{\text{out}} = \mathbf{\Lambda}(\pi - a)$, and $\Delta\mathbf{I} = \mathbf{I}_{\text{in}} - \mathbf{I}_{\text{out}}$.

*Remark* B.2 (Origin of the $-\Delta\mathbf{I}$ term in Equation (B.8)). A reader may expect the matching condition for a second-order ODE to be simply $\mathbf{u}$ continuous at $y = a$. The $-\Delta\mathbf{I}$ term arises because the particular solution $\mathbf{u}^*(y) = \mathbf{M}^{-1}\mathbf{c}(y)$ is piecewise constant and hence discontinuous at $y = a$: it jumps by $\mathbf{u}^*_{\text{in}} - \mathbf{u}^*_{\text{out}} = \mathbf{M}^{-1}\Delta\mathbf{I}$. For the total potential $\mathbf{u}$ to remain physically smooth (as required by the convolution with the $L^1$ kernel), the complementary part $\mathbf{v} = \mathbf{u} - \mathbf{u}^*$ must carry an equal and opposite jump. Writing out the continuity condition for $\mathbf{u}$ at $y = a^-$ and $y = a^+$ and rearranging in terms of the coefficient vectors $\mathbf{a}$ and $\mathbf{c}$ of $\mathbf{v}$ yields Equation (B.8); the term $\mathbf{u}^*_{\text{in}} - \mathbf{u}^*_{\text{out}} - \Delta\mathbf{I} = \mathbf{M}^{-1}\Delta\mathbf{I} - \Delta\mathbf{I} = (\mathbf{M}^{-1} - \mathbf{I})\Delta\mathbf{I}$ captures this compensation. Equation (B.9) follows from the standard condition that $\partial_y \mathbf{u}$ is continuous at $y = a$ (the particular solutions are constant, so $\partial_y \mathbf{u}^* = \mathbf{0}$ on each arc and only the complementary derivatives enter).

#### B.5 Closed-Form Coefficient Vectors

Solving Equations (B.8) and (B.9) yields

$$\mathbf{a} = -\mathbf{K}^{-1}\big(\mathbf{u}^*_{\text{in}} - \mathbf{u}^*_{\text{out}} - \Delta\mathbf{I}\big), \tag{B.10}$$

$$\mathbf{c} = -\sinh^{-1}\!\big(\mathbf{\Lambda}_{\text{out}}\big)\sinh\!\big(\mathbf{\Lambda}_a\big)\,\mathbf{a}, \tag{B.11}$$

where

$$\mathbf{K} \;=\; \cosh\!\big(\mathbf{\Lambda}_a\big) + \cosh\!\big(\mathbf{\Lambda}_{\text{out}}\big)\sinh^{-1}\!\big(\mathbf{\Lambda}_{\text{out}}\big)\sinh\!\big(\mathbf{\Lambda}_a\big). \tag{B.12}$$

Substituting back into Equation (B.7) gives the complete closed-form steady-state profile:

$$\boxed{\mathbf{u}(y) = \begin{cases} \mathbf{u}^*_{\text{in}} + \cosh\!\big(\mathbf{\Lambda}y\big)\,\mathbf{a}, & |y| \le a, \\[2ex] \mathbf{u}^*_{\text{out}} - \cosh\!\big(\mathbf{\Lambda}(\pi - |y|)\big)\,\sinh^{-1}\!\big(\mathbf{\Lambda}_{\text{out}}\big)\sinh\!\big(\mathbf{\Lambda}_a\big)\,\mathbf{a}, & a < |y| \le \pi. \end{cases}} \tag{B.13}$$

The firing-rate profiles follow by threshold rectification.

### C Power Spectral Density, Bandwidth, and Spectral Criticality

#### C.1 Stochastic System and Linear Noise Approximation

*Remark* C.1 (Validity of the PSD derivation). The LNA requires only $\alpha < 0$ (stable fixed point) and is valid for any $\Delta$. The exact PSD formula Equation (C.3) is valid for any $\alpha < 0$, whether the system is in the oscillatory ($\Delta < 0$) or overdamped ($\Delta > 0$) regime. When $\Delta > 0$, the eigenvalues $\lambda_{1,2} = \alpha \pm \kappa$ (where $\kappa = \sqrt{\Delta}/(2\tau\tau') > 0$) are real and negative; the denominator in Equation (C.3) reduces to $(\omega^2 + \lambda_1^2)(\omega^2 + \lambda_2^2)$, giving a monotone, peakless spectrum consistent with an overdamped system. The *additional* requirement $\Delta < 0$

is needed only for the Lorentzian approximation Equation (C.4) and the spectral sharpness index $\mathcal{C}$, both of which presuppose a resonance frequency $\omega_0 = \sqrt{-\Delta}/(2\tau\tau')$. In the limit-cycle regime ($\alpha > 0$), the fixed point is unstable and the LNA does not apply; a Floquet-theory treatment is deferred to future work.

State-dependent multiplicative noise (consistent with Poisson-like neural variability) is incorporated via the macroscopic SDEs:

$$\frac{d}{dt}\begin{pmatrix}\psi \\ \phi\end{pmatrix} = \begin{pmatrix}F_\psi(\psi,\phi) \\ F_\phi(\psi,\phi)\end{pmatrix} + \begin{pmatrix}\frac{\sigma_\psi}{\tau}\sqrt{\psi(t)} \\ \frac{\sigma_\phi}{\tau'}\sqrt{\phi(t)}\end{pmatrix} \odot \begin{pmatrix}\epsilon_\psi(t) \\ \epsilon_\phi(t)\end{pmatrix}, \tag{C.1}$$

where $\epsilon_\mu(t)$ are independent standard Gaussian white noise processes. The LNA around the active fixed point $(\psi_*, \phi_*)$ reduces the dynamics to a coupled Ornstein–Uhlenbeck process with diffusion matrix

$$\mathbf{D} = \begin{pmatrix}\sigma_\psi^2\psi_*/\tau^2 & 0 \\ 0 & \sigma_\phi^2\phi_*/\tau'^2\end{pmatrix}. \tag{C.2}$$

#### C.2 Exact Power Spectral Density

The cross-spectral density matrix is $\mathbf{S}(\omega) = \mathbf{R}(\omega)\,\mathbf{D}\,\mathbf{R}^\dagger(\omega)$ with $\mathbf{R}(\omega) = (i\omega\mathbf{I} - \mathbf{A})^{-1}$. Using the factorization $|\det(i\omega\mathbf{I} - \mathbf{A})|^2 = [\alpha^2 + (\omega-\omega_0)^2][\alpha^2 + (\omega+\omega_0)^2]$, the $(1,1)$ element yields the exact PSD of the excitatory firing rate:

$$S_R(\omega) = \frac{\sigma_\psi^2(R_* + \theta)(\omega^2 + \gamma_2^2)/\tau^2 + \sigma_\phi^2(R'_* + \theta')(\mathcal{J}'_{\mathrm{r}})^2/(\tau^2\tau'^2)}{[\alpha^2 + (\omega-\omega_0)^2][\alpha^2 + (\omega+\omega_0)^2]}, \tag{C.3}$$

where $\gamma_2 = (\mathcal{J}'_0 - 1)/\tau'$. Here $R_* = \psi_* - \theta$ and $R'_* = \phi_* - \theta'$ are the mean excitatory and inhibitory firing rates, respectively.

#### C.3 Lorentzian Approximation and Bandwidth

Near the Hopf boundary ($\omega_0 \gg |\alpha|$), the non-resonant factor $\alpha^2 + (\omega + \omega_0)^2 \approx 4\omega_0^2$, so Equation (C.3) reduces to

$$S_R(\nu) \approx \frac{\sigma_{\text{eff}}^2/4\omega_0^2}{\alpha^2 + (2\pi\nu - \omega_0)^2} + \frac{\sigma_{\text{eff}}^2/4\omega_0^2}{\alpha^2 + (2\pi\nu + \omega_0)^2}, \qquad \Delta < 0,\ \omega_0 \gg |\alpha|, \tag{C.4}$$

where $\sigma_{\text{eff}}^2$ is the numerator of Equation (C.3) evaluated at $\omega = \omega_0$. The one-sided physical PSD ($\nu > 0$) is therefore a Lorentzian with FWHM bandwidth:

$$\Delta f_{\text{FWHM}} = \frac{|\alpha|}{\pi} = \frac{1}{\pi}\left|\frac{\tau(\mathcal{J}_0' - 1) + \tau'(\mathcal{J}_0 - 1)}{2\tau\tau'}\right|. \tag{C.5}$$

#### C.4 Spectral Sharpness Index

We define the spectral sharpness index as

$$\mathcal{C} = \frac{\omega_0}{|\alpha|} = \frac{2f_0}{\Delta f_{\text{FWHM}}} = \frac{\sqrt{4\tau\tau'\big[\mathcal{J}_\text{r}'\mathcal{J}_\text{r} - (\mathcal{J}_0 - 1)(\mathcal{J}_0' - 1)\big] - \big[\tau(\mathcal{J}_0' - 1) + \tau'(\mathcal{J}_0 - 1)\big]^2}}{\big|\tau(\mathcal{J}_0' - 1) + \tau'(\mathcal{J}_0 - 1)\big|}, \tag{C.6}$$

which equals twice the standard quality factor $Q = \omega_0/(2|\alpha|)$ and corresponds to the spectral sharpness index $\mathcal{C}$ reported in the main text. As $\mathcal{J}_0 \to \mathcal{J}_0^*$ (the Hopf boundary), the bandwidth scales as:

$$\Delta f_{\text{FWHM}} \approx \frac{|\mathcal{J}_0 - \mathcal{J}_0^*|}{2\pi\tau} \longrightarrow 0, \tag{C.7}$$

providing an experimentally measurable signature of critical slowing down.

#### C.5 Numerical Validation

PSD estimates are obtained from 60 s simulations of the full nonlinear SDE using the Euler–Maruyama scheme ($\Delta t = 10^{-4}$ s), with observed firing rates evaluated via Welch's method (Hann window, $L = 2048$, overlap 1024, $N_{\text{fft}} = 8192$, $f_s = 1$ kHz) to verify the analytical

FWHM predictions.

### D VIM Fitting Procedure

#### D.1 Data

Population-averaged instantaneous firing rates (kernel-smoothed PSTH, 10 ms bins) from $n = 18$ VIM thalamic neurons at 100 Hz and $n = 10$ neurons at 200 Hz DBS (9 neurons recorded at both frequencies) are available at nine frequencies: 1, 2, 3, 5, 10, 20, 30, 100, and 200 Hz.

#### D.2 Objective Function

The mean-field ODE (Equations (A.7) and (A.8)) has 11 free parameters. We minimize a frequency-proportional weighted sum of squared residuals over $t \in [0, 2.5]$ s:

$$\mathcal{L}(\mathbf{p}) = \sum_{k=1}^{9} w_k \int_0^{2.5} \left[ R_{\text{sim}}(t; f_k) - R_{\text{model}}(t; f_k, \mathbf{p}) \right]^2 dt, \tag{D.1}$$

where $w_k = f_k / f_{\text{min}}$ prioritizes clinically relevant high-frequency conditions ($\geq$100 Hz), $R_{\text{sim}}$ is the population-averaged PSTH, and $R_{\text{model}}$ is the model output at stimulation frequency $f_k$.

#### D.3 Optimization

We use `fmincon` (MATLAB R2023b, SQP algorithm, 10,000 function evaluations) with physiologically motivated parameter bounds: $\tau, \tau' \in [0.005, 1.0]$ s, $\mathcal{J}_0 \in [0, 5]$, $\beta \in [-0.01, 0]$ Hz$^{-1}$, $I_0, I_{\text{b}}, I'_{\text{b}}, \theta, \theta'$ bounded by plausible spike-rate scales. The optimizer is run from 20 random restarts to mitigate local minima; the solution with the smallest objective value is retained.

#### D.4 Fitted Parameters

The fitted parameter vector $\hat{\mathbf{p}}$ is stored in `optimized_params_all.mat` (field `p_fit`, $1 \times 11$ double). Confirmed values read directly from the file:

| Parameter | Symbol | Value | Units |
|---|---|---|---|
| Excitatory time constant | $\hat{\tau}$ | 0.0109 | s |
| Inhibitory time constant | $\hat{\tau}'$ | 1.0000 (at bound) | s |
| E→E eff. coupling | $\hat{\mathcal{J}}_0$ | 0.5898 | — |
| I→I eff. coupling | $\hat{\mathcal{J}}_0'$ | $\approx 0$ | — |
| E→I eff. coupling | $\hat{\mathcal{J}}_\mathrm{r}$ | 0.2213 | — |
| I→E eff. coupling | $\hat{\mathcal{J}}_\mathrm{r}'$ | $-5.516$ | — |
| E background input | $\hat{I}_\mathrm{b}$ | 16.84 | a.u. |
| I background input | $\hat{I}_\mathrm{b}'$ | 0.0018 | a.u. |
| Pulse amplitude | $\hat{I}_0$ | $1.846 \times 10^4$ | a.u. |
| I coupling to DBS | $\hat{\eta}$ | 0.03570 | — |
| Synaptic depression coeff. | $\hat{\beta}$ | $-0.003854$ | $\mathrm{Hz}^{-1}$ |

The fact that $\hat{\tau}'$ is at its upper bound indicates that the effective inhibitory integration time constant is not well resolved from the excitatory rate data alone. This value should be interpreted as the effective population-level time constant of the inhibitory feedback pathway, which may encompass multiple synaptic stages and integrate over a longer timescale than individual GABAergic cells; population-level effective time constants routinely exceed cellular ones [1]. The near-zero $\hat{\mathcal{J}}_0'$ implies that I→I recurrence is negligible at the fitted operating point. We note also that the E–I labelling is a notational convenience; the coupling signs are unconstrained, and a near-zero $\hat{\mathcal{J}}_0'$ or atypical sign in another weight is admissible in data-driven fits.

#### D.5 LNA Applicability and PSD Comparison

The LNA requires only $\alpha < 0$ (stable fixed point), which is satisfied at the fitted parameters ($\alpha < 0$, confirmed numerically). The exact PSD formula (Equation (C.3)) therefore applies. The fitted $\hat{\tau}' = 1.0$ s yields discriminant $\Delta(\hat{\mathbf{p}}) > 0$ (overdamped, real eigenvalues $\lambda_{1,2} = \alpha \pm \kappa$). By Remark C.1, the denominator of Equation (C.3) evaluates to $(\omega^2 + \lambda_1^2)(\omega^2 + \lambda_2^2)$, giving a monotone, peak-less spectrum. This is a genuine LNA prediction, not a phenomenological overlay: the absence of a resonant peak is consistent with VIM thalamus not exhibiting autonomous beta oscillations. The analytical curve in Fig. 2C of the main text is this overdamped LNA spectrum, scaled to match the empirical peak above the estimated Poisson noise floor. The spectral sharpness index $\mathcal{C}$ requires $\Delta < 0$ and is not applicable to the VIM data; no claim about Hopf proximity is made from this comparison.

### SI Figures

### E Transfer Function: Michaelis–Menten to ReLU

The intrinsic firing rate of each population is described by a Michaelis–Menten (MM) saturation function:

$$R_\psi(u) = \frac{[u-\theta]_+}{1 + [u-\theta]_+/R_{\max}}, \qquad [x]_+ = \max(x, 0), \tag{E.1}$$

where $\theta$ is the spiking threshold and $R_{\max}$ is the maximum sustained firing rate [1, 2]. For inputs well below saturation, $[u-\theta]_+ \ll R_{\max}$, the denominator collapses to unity and Equation (E.1) reduces self-consistently to the threshold-linear (ReLU) form $R_\psi \approx \max(u-\theta, 0)$. This reduction is self-consistent in the sense that if the ReLU fixed point satisfies $R_* \ll R_{\max}$, then the MM correction $(R_*/R_{\max})$ is negligible and ReLU is a valid approximation throughout the dynamics.

For the default parameters (Table S1), single-population firing rates under DBS reach at

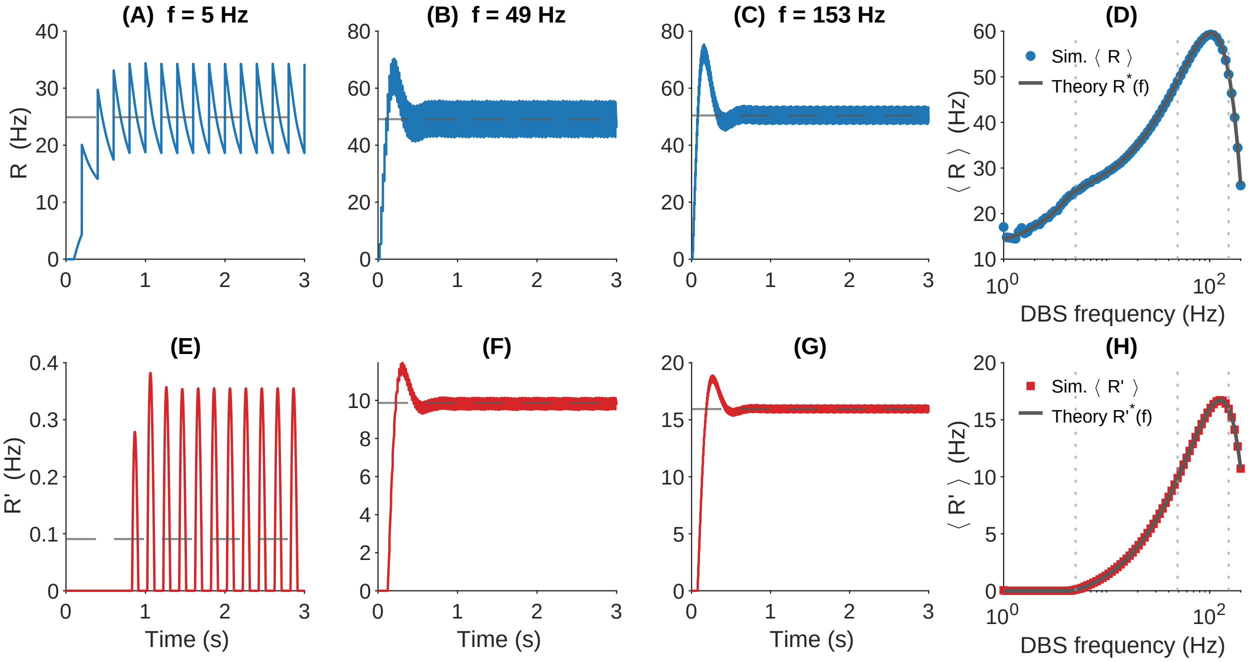


Figure S1: **Fig. S1. DBS-modulated dynamics: spatial simulation versus mean-field theory.** Columns 1–3 show representative time traces at $f \approx 5$, 49, and 153 Hz (the three frequencies on the 80-point log-spaced simulation grid nearest to 5, 50, and 150 Hz, respectively). **(A)–(C)** Excitatory firing rate $R(t)$ at the central neuron (blue); grey dashed: mean-field prediction $R_*(f)$. **(E)–(G)** Inhibitory firing rate $R'(t)$ (red); grey dashed: mean-field prediction $R'^*(f)$. **(D)** Spatial- and time-averaged excitatory rate $\langle R \rangle$ versus DBS frequency (log scale); blue circles: simulation; grey solid: mean-field theory. **(H)** Same for $\langle R' \rangle$. Vertical dotted lines in (D,H) mark the three representative frequencies. Parameters: $N = 100$, $\tau = \tau' = 0.1$ s, $J_0 = 0.015$, $J_0' = -0.006$, $J_\mathrm{r} = 0.012$, $J_\mathrm{r}' = -0.128$, $l = \pi/3$, $\theta = \theta' = 10$, $I_\mathrm{b} = 16$, $I_\mathrm{b}' = 0$, $I_0 = 16000$, $\beta = -0.004$ Hz$^{-1}$, $\eta = 0.04$, $\Delta t = 0.1$ ms.

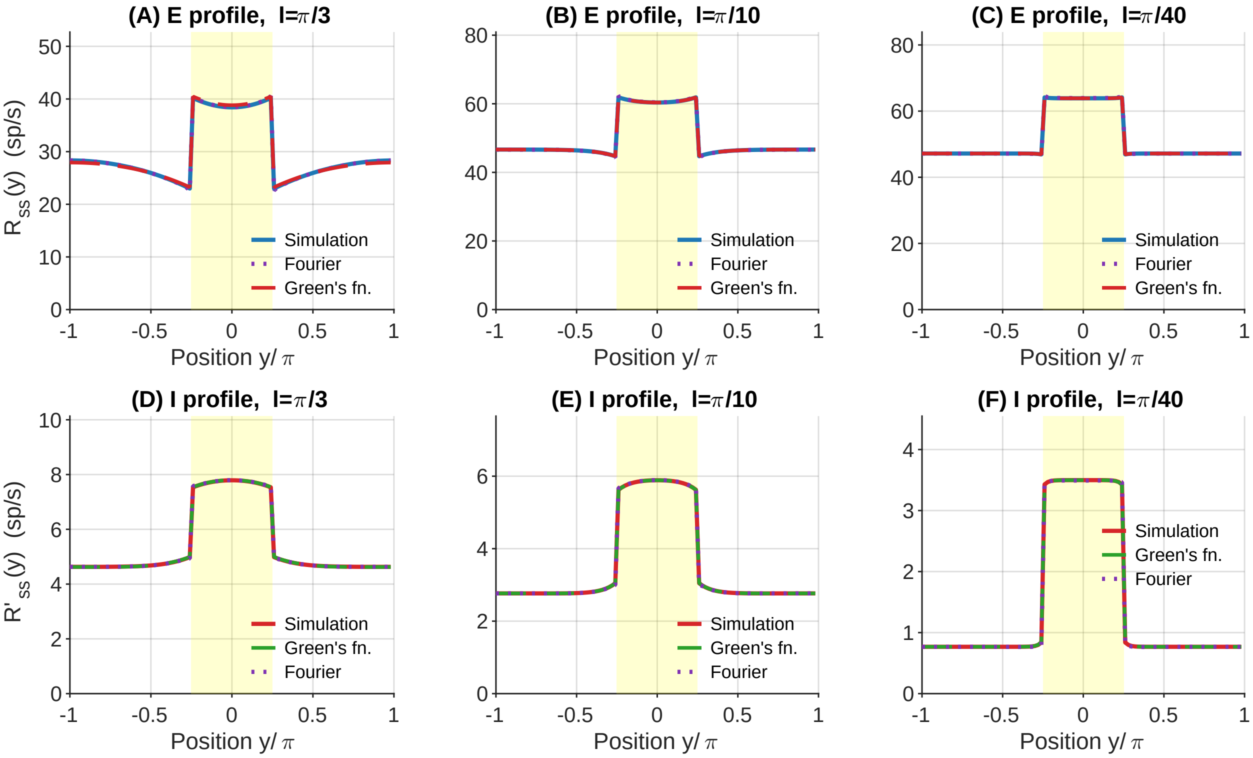


Figure S2: **Fig. S2. Spatial profile accuracy across connectivity decay lengths.** Columns: $l = \pi/3$, $\pi/10$, $\pi/40$. Top row: excitatory profiles **(A)–(C)**; bottom row: inhibitory profiles **(D)–(F)**. In top panels (A)–(C): blue solid = simulation; purple dotted = Fourier (500 modes); red dashed = Green's function. In bottom panels (D)–(F): red solid = simulation; green dashed = Green's function; purple dotted = Fourier. Yellow shading: stimulated region $|y| \leq \pi/4$. Decreasing $l$ produces progressively more localized, sharply demarcated activity bumps. Both analytical methods are consistent with simulation for all three values of $l$, validating the exact periodic kernel coefficient (Equation (B.3)) and the outer-arc boundary condition of the Green's function (SI Appendix, Section B.5). Parameters: $J_0 = 0.0126$, $J_0' = -0.0063$, $J_{\mathrm{r}} = 0.0063$, $J_{\mathrm{r}}' = -0.1885$, $\theta = \theta' = 10$, $I_0 = 2500$, $I_{\mathrm{b}} = 80$, $I_{\mathrm{b}}' = 10$, $\beta = -0.003$ Hz$^{-1}$, $\eta = 0.1$, $\Delta t = 0.1$ ms, $f = 100$ Hz, $d_{\mathrm{stim}} = \pi/2$, $N = 100$.

most 10–50 sp/s, well below $R_{\max} \approx$ 200–400 sp/s for thalamic relay and basal ganglia projection neurons. The ReLU approximation is therefore self-consistent for all parameter regimes analyzed here; the full MM form is required only if $I_0$ or $I_{\mathrm{b}}$ is increased substantially beyond the default values.

The linear efficacy factor $F(f) = 1 + \beta f$ is the first-order Taylor expansion of the Tsodyks–Markram steady-state release probability $(1 + f\tau_{\mathrm{rec}}U)^{-1}$ [3, 4]:

$$\frac{1}{1 + f\tau_{\mathrm{rec}}U} = 1 - U\tau_{\mathrm{rec}}f + O(f^2) \;\equiv\; 1 + \beta f + O(f^2), \qquad \beta = -U\tau_{\mathrm{rec}} < 0. \tag{E.2}$$

For glutamatergic corticosubthalamic and thalamostriatal synapses, $U \approx$ 0.3–0.5 and $\tau_{\mathrm{rec}} \approx$ 5–15 ms [3], giving $\beta \approx -0.002$–$-0.007$ Hz$^{-1}$, bracketing the default value $\beta = -0.003$ Hz$^{-1}$ and the VIM fitted value $\hat{\beta} \approx -0.0038$ Hz$^{-1}$. The first-order approximation is accurate when $f|\beta| \ll 1$ (equivalently $f \ll |\beta|^{-1}$). For the default $|\beta|^{-1} \approx 333$ Hz this condition is well satisfied below 100 Hz; at 200 Hz, where $f|\beta| \approx 0.6$, the linear approximation underestimates $F(f)$ relative to the full Tsodyks–Markram form by roughly 35%. The parameter $\beta$ is therefore best interpreted as a fitted phenomenological slope rather than a direct measurement of $-U\tau_{\mathrm{rec}}$, and predictions at $f \geq 150$ Hz should be treated with corresponding caution.

### F Nucleus-Specific Model Instantiations

The bi-population structure maps onto the four DBS targets as follows. In each case, Population 1 represents the directly stimulated nucleus and Population 2 its primary inhibitory partner within the same circuit loop.

**VIM thalamus.** GABAergic projections from the thalamic reticular nucleus (RT) provide the primary feedforward inhibition onto relay neurons in VIM [5]. Population 2 represents RT neurons; $J_{\phi\phi}$ captures RT self-inhibition and $J_{\phi\psi}$ the VIM→RT excitatory drive. The VIM–RT circuit implements gain modulation rather than threshold suppression under the fitted parameters ($f_{\rm th}$ falls outside the clinical frequency range), consistent with the monotone increase of VIM firing rate with stimulation frequency observed experimentally [6].

**Subthalamic nucleus (STN).** The reciprocal STN–GPe (globus pallidus externus) loop dominates STN dynamics [7, 8]. Population 2 represents GPe neurons; $J_{\phi\phi}$ captures GABAergic GPe self-inhibition and $J_{\phi\psi}$ the STN→GPe glutamatergic drive. The negative $J'_{\rm r} < 0$ (I→E, i.e., GPe→STN) reflects the major GABAergic projection from GPe back to STN. The BEB threshold $f_{\rm th}^{\rm STN} \approx 81.6$ Hz lies below the clinically standard stimulation range of 130–180 Hz [9, 10], consistent with the suppressive regime being operative throughout standard therapy.

**Substantia nigra pars reticulata (SNr).** GABAergic axon collaterals among SNr projection neurons provide robust recurrent inhibition within Population 2 [11, 12, 13]. The SNr threshold $f_{\rm th}^{\rm SNr} \approx 24.7$ Hz is considerably lower than for STN, reflecting the stronger recurrent inhibition ($\mathcal{J}'_0$) fitted from the steeper frequency–response slope observed in [6].

**Thalamic reticular nucleus (RT).** RT is classified here as a gain-modulation nucleus (monotone increasing $R_*(f)$), pairing with VIM in the bifurcation-scaling collapse. The circuit interpretation is that RT receives direct DBS-induced excitation, which is amplified by its own recurrent GABAergic collaterals [5]. The absence of an observable $f_{\rm th}$ in the measured range is reproduced when $\mathcal{J}'_0$ is small (weak self-inhibition) and $\beta$ is only moderately negative.

### G Slow Synaptic Plasticity and Therapeutic Accumulation

The main analysis treats $\mathcal{J}_0$ as time-invariant, capturing acute ($\lesssim 1$ s) effects via the frequency-dependent drive $F(f)$. On the timescale of spike-timing-dependent plasticity ($10^2$–$10^3$ s) [14, 15], sustained high-frequency DBS progressively reduces postsynaptic activity via the BEB, which in turn reduces Hebbian potentiation. A minimal phenomenological rule capturing this feedback is:

$$T_{\mathrm{LTD}}\,\dot{\mathcal{J}}_0 = -\gamma_{\mathrm{LTD}}\,R_*(f)^2 + \gamma_{\mathrm{LTP}}\left(\mathcal{J}_0^{\mathrm{target}} - \mathcal{J}_0\right), \tag{G.1}$$

where the first term is activity-dependent depression proportional to the square of the mean firing rate and the second is homeostatic restoration toward a target coupling. The slow fixed point is:

$$\mathcal{J}_0^{\infty}(f) = \mathcal{J}_0^{\mathrm{target}} - \frac{\gamma_{\mathrm{LTD}}}{\gamma_{\mathrm{LTP}}}\,R_*(f)^2. \tag{G.2}$$

In the pathological state $\mathcal{J}_0$ is assumed to exceed $\mathcal{J}_0^{\mathrm{target}}$. For $f > f_{\mathrm{th}}$, the BEB drives $R_*$ to zero, eliminating the activity-dependent term in Equation (G.1). The homeostatic term then drives $\mathcal{J}_0 \to \mathcal{J}_0^{\mathrm{target}} < \mathcal{J}_0$, progressively moving the operating point away from the Hopf boundary $\mathcal{J}_0^*$. This provides a candidate account of the gradual accumulation of therapeutic benefit observed over days to weeks of continuous DBS [10]: acute suppression via the BEB is augmented by slow homeostatic reduction of pathological recurrent coupling.

*Remark* G.1 (Status and validation path). Equation Equation (G.1) is a phenomenological hypothesis, not a derived prediction. Direct validation requires chronic LFP recordings tracking beta bandwidth $\Delta f_{\mathrm{FWHM}} = |\alpha|/\pi$ over weeks of DBS alongside an independent estimate of $\tau$ from in-vitro synaptic physiology. The acute subsystem (Sections A–C) remains valid at any instantaneous value of $\mathcal{J}_0$, regardless of whether the slow plasticity rule is operative.

### H Inhibitory Halo: Non-Trivial Spatial Solution

The sign of the coefficient vector $\mathbf{a} = -\mathbf{K}^{-1}(\mathbf{u}_{\mathrm{in}}^* - \mathbf{u}_{\mathrm{out}}^* - \Delta\mathbf{I})$ (Section B.5, Equation (B.10)) determines the qualitative character of the spatial profile.

**Activity bump (a < 0).** This is the classical CANN activity bump: $\psi_{\mathrm{ss}}$ is maximized at $y = 0$ and decays smoothly toward the background level. The electrode region is more active than the surround.

**Inhibitory halo (a > 0).** When $|\mathcal{J}_{\mathrm{r}}'|$ is large enough relative to $\mathcal{J}_0$, the inner cosh term elevates $\psi_{\mathrm{ss}}$ toward the electrode edge rather than toward the center. The outer arc then shows a trough below background — an inhibitory collar surrounding the stimulated region. The suppression depth at the electrode edge is:

$$\delta R_{\mathrm{halo}} = \left[\tanh(\mathbf{\Lambda}_{\mathrm{out}})\,\sinh(\mathbf{\Lambda}_a)\,\mathbf{a}\right]_1, \tag{H.1}$$

where subscript 1 denotes the first (excitatory) component.

This annular suppression pattern is a generic feature of lateral-inhibition-mediated surround suppression: a center-surround connectivity profile, in which direct excitatory drive falls off less steeply with distance than recurrent inhibitory drive, naturally produces a ring of net suppression flanking the directly activated region. Within the CANN model, $\delta R_{\mathrm{halo}}$ scales linearly with $I_0$ through $\mathbf{a} \propto \Delta\mathbf{I}$; the spatial width of the halo is governed by $\mathbf{\Lambda} = \mathbf{M}^{1/2}/l$ and is independent of stimulation amplitude. Whether the inhibitory collar is experimentally detectable under DBS-scale currents requires recordings from neurons at known distances from the electrode while co-varying stimulation amplitude.

### References


[1] Peter Dayan and L. F. Abbott. *Theoretical Neuroscience: Computational and Mathematical Modeling of Neural Systems.* MIT Press, Cambridge, MA, 2001.

[2] Wulfram Gerstner, Werner M. Kistler, Richard Naud, and Liam Paninski. *Neuronal Dynamics: From Single Neurons to Networks and Models of Cognition.* Cambridge University Press, Cambridge, UK, 2014.

[3] Misha V. Tsodyks and Henry Markram. The neural code between neocortical pyramidal neurons depends on neurotransmitter release probability. *Proceedings of the National Academy of Sciences*, 94(2):719–723, 1997. doi: 10.1073/pnas.94.2.719.

[4] L. F. Abbott, J. A. Varela, Kamal Sen, and Sacha B. Nelson. Synaptic depression and cortical gain control. *Science*, 275(5297):220–224, 1997. doi: 10.1126/science.275.5297.220.

[5] Didier Pinault. The thalamic reticular nucleus: Structure, function and concept. *Brain Research Reviews*, 46(1):1–31, 2004. doi: 10.1016/j.brainresrev.2004.04.008.

[6] Luka Milosevic, Suneil K. Kalia, Mojgan Hodaie, Andres M. Lozano, Milos R. Popovic, William D. Hutchison, and Milad Lankarany. A theoretical framework for the site-specific and frequency-dependent neuronal effects of deep brain stimulation. *Brain Stimulation*, 14(4):807–821, 2021. doi: 10.1016/j.brs.2021.04.022.

[7] Jonathan E. Rubin and David Terman. High frequency stimulation of the subthalamic nucleus eliminates pathological thalamic rhythmicity in a computational model. *Journal of Computational Neuroscience*, 16(3):211–235, 2004. doi: 10.1023/B:JCNS.0000025686.47117.67.

[8] Atsushi Nambu, Hironobu Tokuno, and Masahiko Takada. Functional significance of the cortico-subthalamo-pallidal "hyperdirect" pathway. *Neuroscience Research*, 43(2):111–117, 2002. doi: 10.1016/s0168-0102(02)00027-5.

[9] Cameron C. McIntyre, Marc Savasta, Lydia Kerkerian-Le Goff, and Jerrold L. Vitek. Uncovering the mechanism(s) of action of deep brain stimulation: activation, inhibition, or disruption? *Clinical Neurophysiology*, 115(6):1239–1248, 2004. doi: 10.1016/j.clinph.2003.12.024.

[10] Wolf-Julian Neumann, Leon A. Steiner, and Luka Milosevic. Neurophysiological mechanisms of deep brain stimulation across spatiotemporal resolutions. *Brain*, 146(11):

4456–4468, 2023. doi: 10.1093/brain/awad239.

[11] J. M. Deniau, C. Hammond, A. Riszk, and J. Feger. Neuronal interactions in the substantia nigra pars reticulata through axon collaterals of the projection neurons. *Experimental Brain Research*, 47(1):105–113, 1982. doi: 10.1007/BF00235891.

[12] Philippe Mailly, Stéphane Charpier, Anne Menetrey, and Jean-Michel Deniau. Three-dimensional organization of the recurrent axon collateral network of the substantia nigra pars reticulata neurons in the rat. *Journal of Neuroscience*, 23(12):5247–5257, 2003. doi: 10.1523/JNEUROSCI.23-12-05247.2003.

[13] Matthew H. Higgs and Charles J. Wilson. The inhibitory microcircuit of the substantia nigra provides feedback gain control of the basal ganglia output. *eLife*, 3:e02397, 2014. doi: 10.7554/eLife.02397.

[14] Guo-qiang Bi and Mu-ming Poo. Synaptic modifications in cultured hippocampal neurons: dependence on spike timing, synaptic strength, and postsynaptic cell type. *Journal of Neuroscience*, 18(24):10464–10472, 1998. doi: 10.1523/JNEUROSCI.18-24-10464.1998.

[15] Daniel E. Feldman. The spike-timing dependence of plasticity. *Neuron*, 75(4):556–571, 2012. doi: 10.1016/j.neuron.2012.08.001.
